# *In vivo* HSPC gene therapy of hemoglobinopathies without drug selection of corrected cells

**DOI:** 10.64898/2026.09.15.751289

**Authors:** Akshara Sakunthala Velmurugan, Kiriaki Paschoudi, Jack A. Queenan, Sucheol Gil, Aidan Maynard, Charlie E. Fontana, Reva Sharma, Anika Long, Chang Li, Hongjie Wang, Anna K. Anderson, Aphrodite Georgakopoulou, Nikoletta Psatha, Maria Giannaki, Efthymia Vlachaki, Nouraiz Ahmed, Alexander A Sousa, David R. Liu, Evangelia Yannaki, André Lieber, Karthik V Karuppusamy

## Abstract

*In vivo* hematopoietic stem/progenitor cell (HSPC) gene therapy remains limited by low gene-editing efficiency and a lack of clinically applicable selection strategies to enrich therapeutically corrected progeny. We used *in vivo* base and prime editing to introduce a nonpathogenic EPOR variant into HSPCs, conferring erythropoietin hypersensitivity and promoting preferential expansion of gene-corrected erythroid cells. EPOR editing was combined with three therapeutic approaches for the correction of hemoglobinopathies: γ-globin gene addition, γ-globin reactivation, or correction of the sickle cell disease mutation. Tropism-modified helper-dependent adenoviral vectors (HDAd6/3+) targeting HSPCs were used to simultaneously deliver the EPOR-editing machinery and the corresponding therapeutic components. *In vitro* studies in an erythroid progenitor cell line and primary CD34^+^ cells demonstrated that the EPOR^W439*^ variant conferred a strong proliferative advantage to therapeutically modified erythroid progenitors. Mice humanized with CD34^+^ cells from a β^0^/β^0^-thalassemia patient were subjected to EPOR^W439*^-mediated EPO hypersensitivity alongside a therapeutic γ-globin transgene which resulted in >70% HbF-positive erythroid cells and substantial reversion of the disease-associated phenotype, including reduced oxidative stress, near-complete elimination of splenic iron deposition, and reduced splenomegaly. Importantly, these effects were achieved after simple intravenous administration of the vectors following HSPC mobilization and cytokine prophylaxis, without subsequent pharmacologic selection. Together, these findings establish a strategy to amplify the therapeutic benefit of otherwise limited *in vivo* HSPC gene editing for hemoglobinopathies.

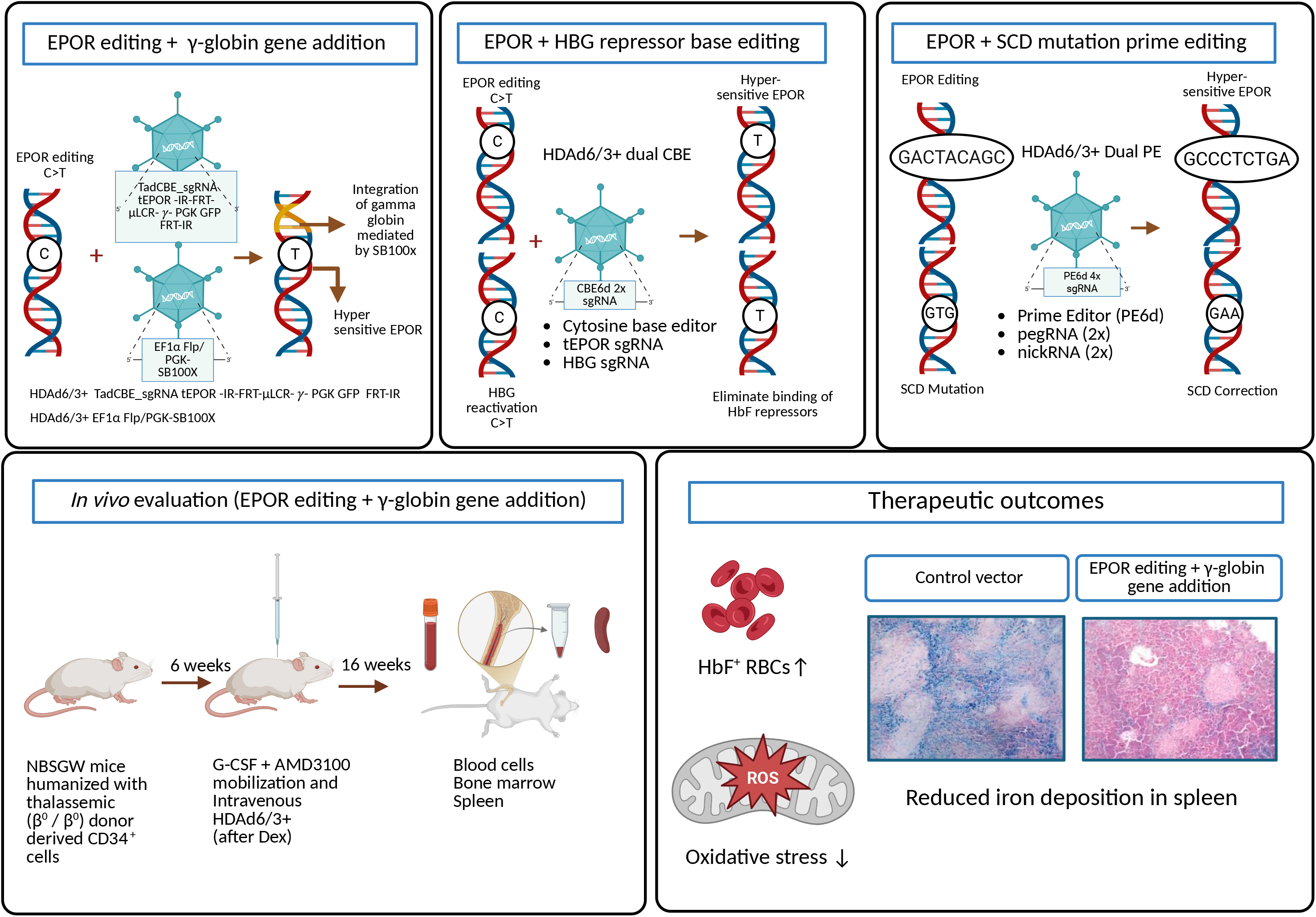

**Key Points:**

- EPOR editing confers a selective growth advantage to engineered erythroid cells under low-EPO conditions.
- In vivo EPOR editing enriches therapeutic erythroid output and supports a chemotherapy-free gene therapy strategy for hemoglobinopathies.

## Introduction

*Ex vivo* hematopoietic stem and progenitor cell (HSPC) gene therapies for hemoglobinopathies are costly and complex(1). We have developed an *in vivo* approach that is now being evaluated in a clinical trial for X-linked chronic granulomatous disease (X-CGD) (https://clinicaltrials.gov/study/NCT06876363). In this approach, HSPCs are mobilized and engineered *in vivo* using intravenously administered, HSC-tropic helper-dependent adenovirus (HDAd) vectors(2). A fraction of engineered HSPCs migrate to the bone marrow and contribute to multilineage hematopoiesis; however, the frequency of long-term gene-modified HSPCs is insufficient for therapy of hemoglobinopathies. The current approach therefore requires chemotherapy-based enrichment either by O^6^BG/BCNU or O^6^BG/Temozolomide(3–5). This chemo-selection strategy can cause significant nonhematologic toxicities, including gastrointestinal and pulmonary complications, and its use may be further complicated in patients with hemoglobinopathies receiving myelosuppressive therapies such as hydroxyurea(6–8). Developing chemotherapy-free enrichment strategies is therefore critical for advancing *in vivo* HSPC gene therapy. Alternative selection strategies using engineered epitopes in CD45 or CD117 have been developed to enable targeted depletion with monoclonal antibodies, antibody-drug conjugates, or CAR T cells(9–12). These approaches may also target non-hematopoietic cell types that express the receptor(13). Consequently, an intrinsic mechanism that selectively expands therapeutically corrected erythroid progeny without altering the non-erythroid hematopoietic compartment could offer a potential alternative. The erythropoietin receptor (EPOR) provides such a potential mechanism. C-terminal EPOR truncations remove negative regulatory domain and prolong EPO-dependent JAK2/STAT5 signaling, resulting in erythropoietin (EPO) hypersensitivity(14,15). Germline EPOR-truncating variants cause primary familial and congenital polycythemia (PFCP), an autosomal-dominant erythrocytosis generally associated with a mild clinical phenotype characterized by increased red blood cell production and low circulating EPO levels(16). The best-characterized example is the c.1316G>A; p.W439* Mäntyranta variant, which removes the C-terminal 70 amino acids of EPOR (17). Importantly, long-term observations of families with germline EPOR-truncating variants have not identified an apparent predisposition to hematologic malignancy or other cancers. This contrasts with somatic EPOR rearrangements described in a subset of high-risk acute lymphoblastic leukemias, which arise in a distinct oncogenic context(18).

C-terminally truncated EPOR variants confer EPO hypersensitivity and enhance erythroid output. However, prior lentiviral and Cas9/HDR approaches require *ex vivo* manipulation and carry risks of random integration or DNA double-strand break (DSB)-associated InDels. (19). Base and prime editing offer a more precise strategy for targeted EPOR modification and *in vivo* selection of edited RBCs. Here, we asked whether this biological advantage could be generated directly *in vivo* and coupled to therapeutic editing for hemoglobinopathies. Using HSPC-tropic HDAd vectors we combined EPOR editing with three therapeutic strategies: γ-globin gene addition, γ-globin reactivation by dual base editing, and correction of the sickle cell disease (SCD) mutation in the *HBB* gene by dual prime editing. Studies were performed in HUDEP-2 cells (human umbilical cord-derived erythroid progenitor cells), primary CD34^+^ cells from healthy donors and patients with β-thalassemia and sickle cell disease (SCD) as well as in humanized mouse models without administration of additional enrichment agents.

## Methods

### Cells

Frozen aliquots of healthy donor-derived mobilized peripheral blood CD34^+^ cells were obtained from the Fred Hutchinson Cancer Center, Seattle, Washington. CD34^+^ cells from SCD patients homozygous for HbS/HbS (N = 3) were immunomagnetically isolated from exchange transfusion blood samples at the George Papanikolaou Hospital, Thessaloniki, Greece. CD34^+^ cells from patients with β-thalassemia major (β⁰/β⁰) (N=3) were collected during two mobilization clinical trials conducted at G. Papanikolaou Hospital(20,21). (NCT00336362 and NCT01206075). All patient-derived samples were obtained after written informed consent in accordance with protocols approved by the appropriate institutional ethics committees and in accordance with the declaration of Helsinki.

### HDAd vectors

HDAd6/3+ is derived from serotype Ad6(22), with the Ad6 fiber knob replaced by an affinity-enhanced Ad3 fiber knob (23).

### Mice

Humanized mouse models (6-8 weeks old; male and female mice were used for this study) were generated by transplanting human CD34^+^ cells from healthy donors or a β-thalassemia donor (β⁰/β⁰) into severely immunodeficient mice (1 × 10⁶ cells/recipient) following partial myeloablation with busulfan (12.5 mg/kg, i.p.). Six weeks after transplantation, mice underwent a GCSF/AMD3100 based 7-day mobilization regimen and received intravenous HDAd injection at a total dose of 8 × 10¹⁰ viral particles. For cytokine prophylaxis, dexamethasone (10 mg/kg, i.p.) was administered 16 and 2 hours before HDAd vector injection(24).

Detailed methods are described in the *Supplemental Information*.

## Results

### Base editing to create naturally occurring truncated EPOR variants in HUDEP-2 cells transduced with lentiviral vectors

We first screened C-terminal truncating EPOR variants that could confer a selective erythroid advantage and assessed whether cytosine or adenine base editing could generate them. Based on target-site compatibility and the desired editing outcomes, we selected cytosine base editing to introduce premature stop codons in the C-terminal region of EPOR. HUDEP-2 cells (EPO-dependent human erythroid progenitor cell line) stably expressing Cas9 or TadCBE were generated by lentiviral transduction followed by puromycin selection (Figs. S1A and S2A). We then transduced these cells with lentiviral vectors co-expressing EPOR-specific sgRNAs and GFP to track transduced cells. The sgRNAs were designed to create InDels around W439 region as well as introduce premature stop codons at Q434, W439, Q474, or Q477, generating a series of C-terminally truncated EPOR variants, including the naturally occurring Mäntyranta variant (EPOR^W439*^)(25). These truncations removed the C-terminal negative regulatory domain within the EPOR cytoplasmic region that attenuates receptor signaling (Figs. S2B-F). We used a gRNA targeting CCR5(R225) as a negative control. Transduced cells were cultured under standard (3 U/mL) or low (0.005 U/mL) EPO concentrations to assess EPO hypersensitivity. EPOR^Q434*^ and EPOR^W439*^ showed significantly greater expansion than the other variants and unedited cells (Figs. S1D,E,G,H), accompanied by a progressive increase in editing frequencies during culture at both EPO concentrations tested (Figs. S1F,I). Among these two variants, the EPOR^W439*^ sgRNA achieved precise target editing without bystander edits observed in the EPOR Q434* sgRNA (Fig. S3A). We further compared two sgRNAs targeting W439* with target cytosines at protospacer positions 5 and 6 (sgRNA-1; 70-1) or positions 4 and 5 (sgRNA-2; 70-2). sgRNA-2 achieved 29% editing compared with 16% for sgRNA-1, representing an approximately 1.8-fold increase in editing efficiency, and was therefore selected for subsequent studies (Figs. S3B-C).

### HDAd vectors mediated base editing to recreate truncated EPOR and integration of the γ-globin gene in HUDEP-2 cells

Our goal was to develop dual-function HDAd vectors, combining EPOR editing to generate the hypersensitive EPOR^Q434*^ or EPOR^W439*^ variants with a therapeutic module for treatment of hemoglobinopathies (Fig. 1A). The therapeutic unit was composed of an integrating γ-globin gene-expressing cassette in which the γ-globin gene was placed under the control of β-globin mini-locus control region (mLCR) and β-globin promoter for erythroid-specific expression. The integrating LCR-γ-globin-GFP cassette is flanked by Inverted Repeats (IRs) for random integration by *Sleeping Beauty* SB100x-transposase, which is provided by a second vector (HDAd.SB100x). The selection unit consists of a cytosine base editor and EPOR-sgRNA, is located outside the IR-flanked transposon, and is therefore expected to be expressed transiently from episomal HDAd genomes.

**Figure 1.**
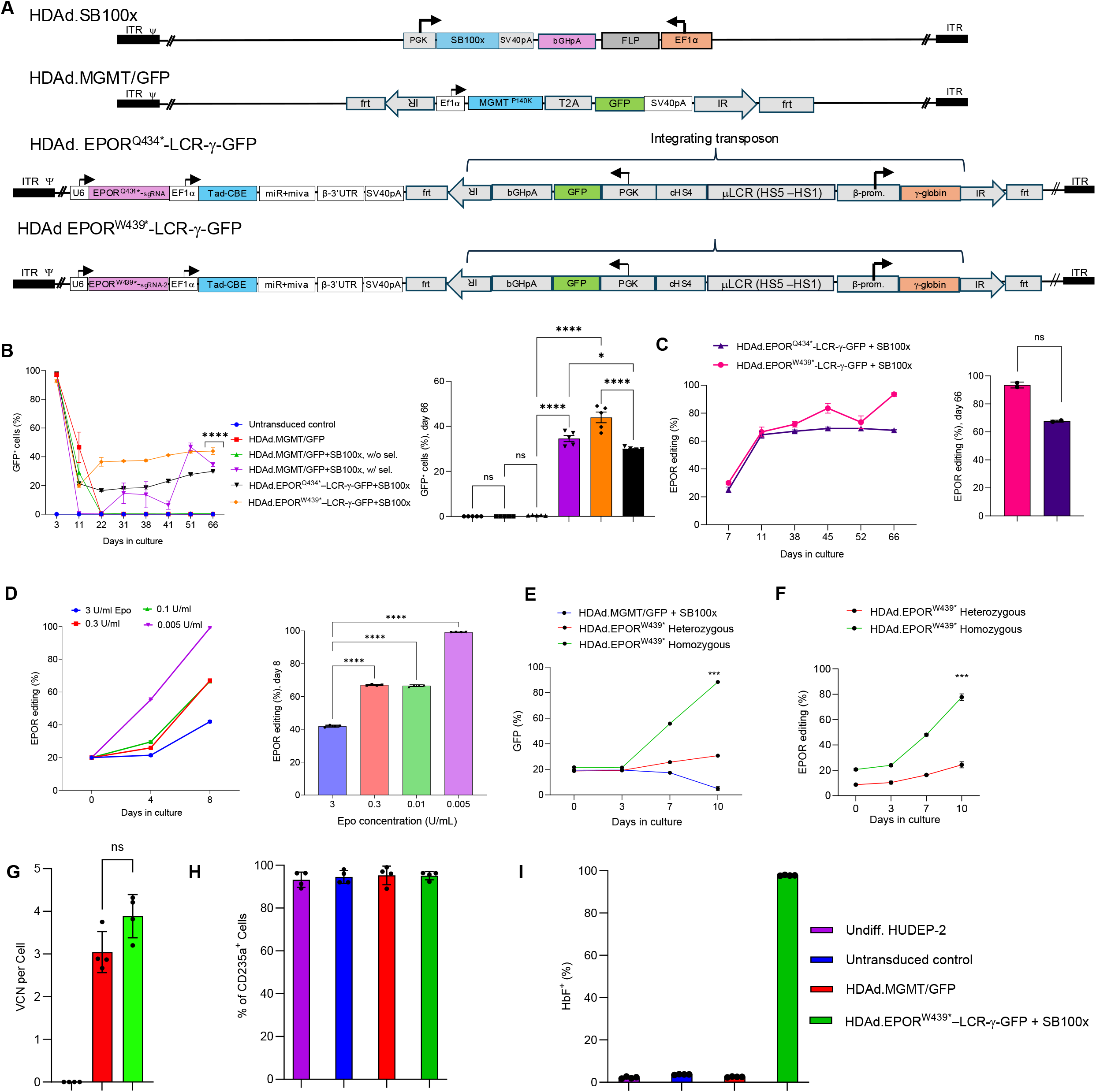
HDAd-mediated EPOR base editing combined with γ-globin gene addition promotes preferential erythroid expansion and HbF expression in HUDEP-2 cells. (A) Schematic representation of the helper-dependent adenoviral (HDAd) vectors used in this study. HDAd.SB100x expresses SB100x transposase and FLP recombinase. HDAd.MGMT/GFP contains an integrating MGMT^P140K^–T2A–GFP transposon cassette. HDAd.EPOR^Q434*^–LCR-γ-GFP and HDAd.EPOR^W439*^–LCR-γ-GFP encode the indicated EPOR base-editing components together with an integrating cassette containing GFP and a β-globin locus control region (LCR)-regulated γ-globin expression cassette. ITR, adenoviral inverted terminal repeat; IR, *Sleeping Beauty* transposase inverted repeat; FRT, FLP recombination target site; LCR, locus control region. **(B)** Longitudinal analysis of GFP-positive HUDEP-2 cells following transduction with the indicated HDAd vectors, with or without SB100x-mediated integration and chemoselection. Right: Percentage of GFP-positive cells on day 66. HDAd.MGMT/GFP+SB100x w/ sel: Cells were subjected to O^6^-BG/BCNU treatment on day 22 to enrich stable integrated cells as per the protocol mentioned in the supplementary methods. Cells were transduced at 2000 MOI/cell. Data represent *n* = 4 independent biological replicates. Endpoint comparisons were analyzed by one-way ANOVA with Šídák’s multiple-comparisons test. **(C)** Longitudinal EPOR-editing frequencies in HUDEP-2 cells transduced with HDAd.EPOR^Q434*^–LCR-γ-GFP or HDAd.EPOR^W439*^–LCR-γ-GFP together with HDAd.SB100x. Right, EPOR-editing frequencies on day 66. Data represent *n* = 2 independent biological replicates; endpoint comparison was performed using an unpaired two-tailed *t* test. **(D)** Enrichment of EPOR^W439*-^edited cells under decreasing EPO concentrations. Cells were cultured in medium containing 3, 0.3, 0.1, or 0.005 U/mL EPO, and EPOR-editing frequencies were measured on days 0, 4, and 8. Right, EPOR-editing frequencies on day 8. Data represents *n* = 4 independent biological replicates; endpoint comparisons were analyzed by one-way ANOVA with Šídák’s multiple-comparisons test. **(E-F)** Preferential enrichment of heterozygous and homozygous EPOR^W439*^-edited cells under low-EPO conditions. Heterozygous or homozygous EPOR^W439*^-edited HUDEP-2 cells were mixed with unedited HUDEP-2 cells at an initial 20:80 ratio and cultured in 0.005 U/mL EPO. Enrichment was monitored by **(E)** GFP positivity and **(F)** EPOR-editing frequency. Data represent *n* = 3 independent biological replicates and were analyzed by two-way ANOVA with Šídák’s multiple-comparisons test. **(G)** Vector copy number (VCN) per cell in the indicated experimental groups, determined by digital PCR. Data represent *n* = 4 independent biological replicates and were analyzed by one-way ANOVA with Šídák’s multiple-comparisons test. **(H)** Percentage of CD235a+ cells on day 11 after erythroid differentiation (**I)** Percentage of HbF-positive cells following erythroid differentiation (ED), determined by flow cytometry. Undifferentiated HUDEP-2 cells, untransduced differentiated cells, HDAd.MGMT/GFP-transduced cells, and cells transduced with HDAd.EPOR^W439*^–LCR-γ-GFP plus HDAd.SB100x were analyzed. For the EPOR^W439*^ group, cells cultured in 0.005 U/mL EPO in panel D were collected on day 8 and subjected to erythroid differentiation. Data represent *n* = 4 independent biological replicates and were analyzed by one-way ANOVA with Šídák’s multiple-comparisons test. Data are presented as mean ± SEM. ns, not significant; *P* < .05; \**P* < .01; \*\**P* < .001; \*\*\**P* < .0001.

HUDEP-2 cells were co-transduced with HDAd.EPOR^Q434*^LCR-γ-GFP or HDAd.EPOR^W439*^-LCR-γ-GFP together with HDAd.SB100x, and GFP marking was monitored longitudinally (Fig. 1B). On day 3, ∼90-100% GFP expression, predominantly from episomal vector genomes, confirmed efficient transduction. Cells were then passaged for 2 months at 3 U/ml of EPO. We compared EPOR-mediated expansion with O^6^BG/BCNU-mediated selection using HDAd.MGMT/GFP + HDAd.SB100x. In control groups, lacking EPOR-mediated expansion or O^6^BG/BCNU selection, GFP marking declined to background levels by day 22, consistent with loss of episomal HDAd genomes and inefficient stable integration. In contrast, after initial decline, GFP marking in the EPOR^W439*^ group increased from 19.86% on day 11 to ∼36.5% by day 22, whereas EPOR^Q434*^ remained at 16% (Fig. S1D). By day 66, GFP marking reached 43.94% for EPOR^W439*^, whereas 29.92% for EPOR^Q434*^. O^6^BG/BCNU selection also enriched MGMT^P140K^-modified cells, but it was accompanied by ∼99% cell loss following selection (data not shown). Notably, EPOR^W439*^ marking exceeded that achieved with the mgmt^P140K^-O^6^BG/BCNU selection system.

Targeted high-throughput sequencing (HTS) analysis showed corresponding enrichment of EPOR-edited alleles, reaching 62% and 95% for HDAd.EPOR^W434*^-LCR-γ-GFP and HDAd.EPOR^W439*^-LCR-γ-GFP editing, respectively (Fig. 1C). We focused subsequent studies on EPOR^W439*^. As observed in lentiviral screening experiments, lowering EPO concentrations progressively enriched EPOR^W439*^-edited cells, with maximal enrichment at 0.005 U/mL EPO (Fig. 1D). This inverse relationship likely reflects increased EPO sensitivity of EPOR^W439-^edited cells, resulting in stronger selective enrichment at lower EPO concentrations. Notably, editing frequencies exceeded GFP positivity, suggesting EPOR editing can occur without stable γ-globin/GFP integrations. Furthermore, we generated HUDEP-2 clones carrying heterozygous-monoallelic or homozygous-biallelic EPOR^W439*^ edits. Only homozygous EPOR^W439*^ cells showed marked enrichment based on GFP marking and editing frequency (Figs. 1E-F), indicating that biallelic editing is required for robust expansion under these conditions. Comparable vector copy numbers excluded differences in transgene integration as a confounding factor (Fig. 1G). Following erythroid differentiation (ED), EPOR^W439*^–LCR-γ-GFP-treated cells were 100% positive for γ−globin (Fig. 1H-I).

Together, these findings demonstrate that HDAd-mediated EPOR^W439*^ cytosine base editing confers EPO hypersensitivity and preferential expansion of erythroid progenitors carrying an integrated therapeutic γ-globin cassette.

### Preferential proliferation of erythroid progenitors derived from EPOR edited CD34^+^ cells from healthy donors

Human CD34^+^ cells, a cell fraction that is enriched for HSPCs, were transduced with integrating HDAd.MGMT/GFP or HDAd.EPOR^W439*^-LCR-γ-GFP vectors and subjected to colony-forming unit (CFU) assays or ED in liquid culture (Fig. 2A). HDAd vectors possessing Ad6/3+ capsids transduced CD34^+^ cells more efficiently than HDAd5/35++ vectors (Fig. 2B), which is consistent with previous observations(22). We therefore used the HDAd6/3+ vector platform in all subsequent studies.

**Figure 2.**
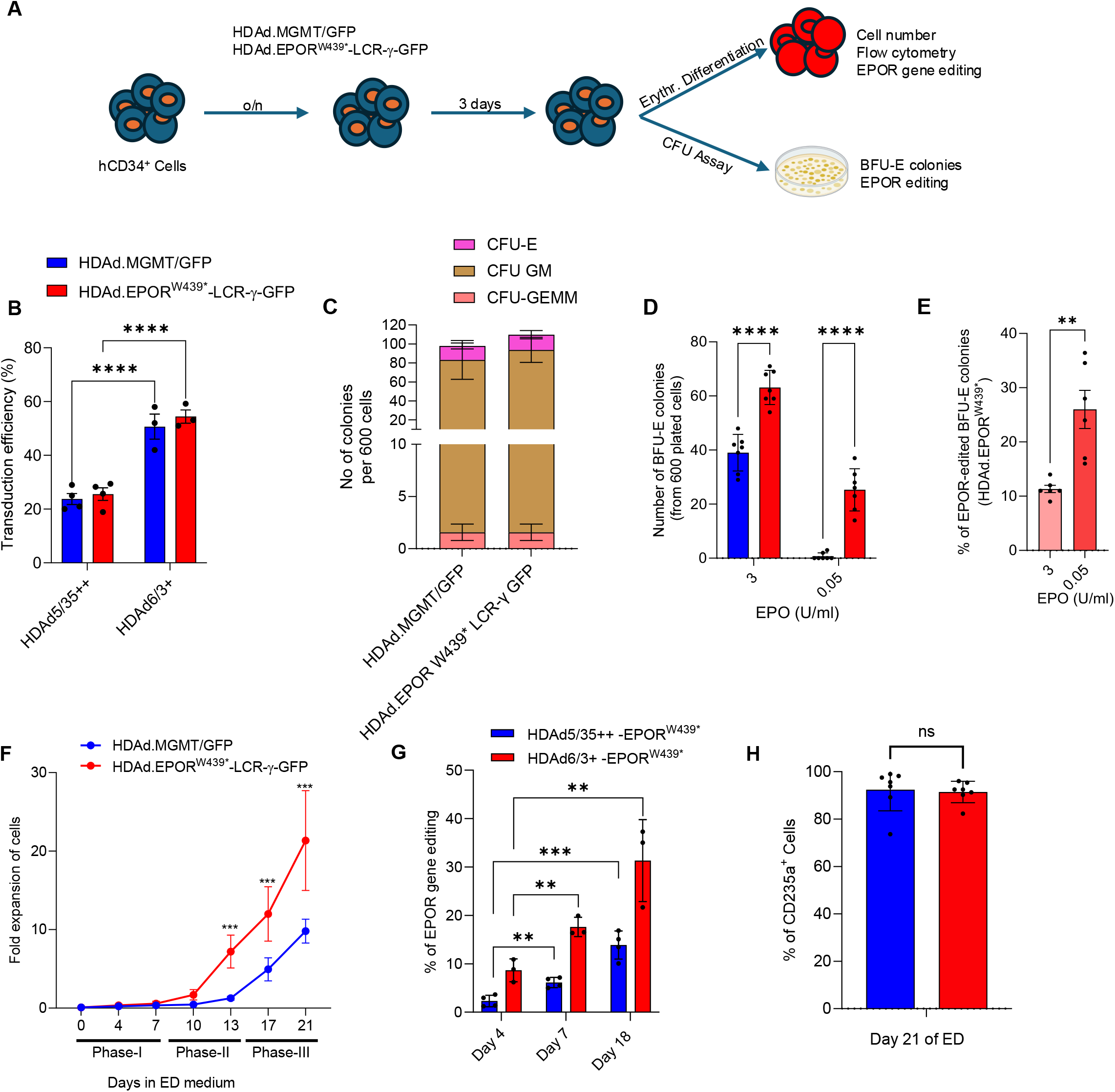
EPOR^W439*^ base editing enhances erythroid output from primary human CD34⁺ cells. **(A)** Experimental schematic. Human CD34⁺ cells, a fraction enriched for hematopoietic stem/progenitor cells (HSPCs), were transduced overnight with HDAd vectors encoding either control (HDAd.MGMT/GFP) or EPOR^W439*^ base editor with γ-globin/GFP cassette (HDAd.EPOR^W439*^-LCR-γ-GFP), followed by 3 days of recovery. Cells were then subjected to erythroid differentiation or colony-forming unit (CFU) assays. Readouts included an *in vitro* colony formation assay, cell counts, EPOR editing, and flow cytometry-based erythroid analysis. **(B)** Transduction efficiency in CD34⁺ cells using HDAd vectors. We observed comparable GFP marking frequencies between control and HDAd.EPOR^W439*^ vectors across different vector doses. HDAd vectors either possessed CD46-targeting HDAd5/35++ capsids or DSG2-targeting HDAd6/3+ capsids. All subsequent experiments were performed with HDAd6/3+ vectors. **(C)** Distribution of CFU-E, CFU-GM, and CFU-GEMM colonies generated from control and EPOR^W439*^-edited CD34^+^ cells. **(D)** BFU-E colony formation under regular (3 U/mL) and low (0.05 U/mL) EPO conditions. EPOR^W439*-^edited cells show significantly more colony numbers compared to control, particularly under low EPO conditions. **(E)** Percentage of EPOR-edited BFU-E colonies. A higher fraction of colonies carry EPOR edits under low EPO conditions, consistent with selective expansion of hypersensitive cells. **(F)** Fold expansion of erythroid cells during differentiation. EPOR^W439*^-edited cells exhibit markedly enhanced proliferation over time compared to control cells. **(G)** Longitudinal EPOR editing frequencies in CD34^+^ cells and differentiated erythroid cells. **(H)** Editing levels increase over time, with higher enrichment observed in EPOR^W439*^ groups. Percentage of CD235a^+^ cells on day 21 of erythroid differentiation, showing comparable erythroid differentiation between control and EPOR^W439*^-edited cultures. Data are presented as mean ± SEM. Statistical comparisons in panels B and D were performed using two-way ANOVA with Šídák’s multiple-comparisons test. Longitudinal analyses in panels F and G were performed using two-way repeated-measures ANOVA with Šídák’s multiple-comparisons test. Comparisons in panels E and H were performed using two-tailed paired *t* tests. ns, not significant; \**P* < .01; \*\**P* < .001; \**P* < .0001.

Previous work showed increased BFU-E formation from individuals carrying EPOR mutations under low Epo conditions(26). We confirmed this in CFU assays with HDAd.EPOR^W439*^-LCR-γ-GFP transduced CD34^+^ cells. Overall CFU composition was comparable between EPOR^W439*^-edited and control cells (Fig. 2C), suggesting that EPOR editing did not affect multilineage differentiation. At 3 U/mL EPO, EPOR^W439*^-edited cells generated more than twice as many burst-forming unit-erythroid (BFU-E) colonies as controls, with a greater advantage for 0.05 U/mL EPO (Fig. 2D). Approximately 10% and 25% of BFU-E colonies carried the desired EPOR^W439*^ edit at 3 and 0.05 U/mL EPO, respectively, demonstrating preferential enrichment under low-EPO conditions (Fig. 2E). These findings were confirmed using a three-phase erythroid differentiation protocol(27). EPOR^W439*^-edited cells showed modest increased expansion during phases I and II, followed by marked expansion during the maturation phase III (Fig. 2F). EPOR^W439*^ editing also progressively increased during differentiation, confirming preferential enrichment of edited erythroid cells (Fig. 2G). Both groups reached comparable frequencies of CD235a^+^ (glycophorin A, a well-established marker of erythroid differentiation) cells by day 21, indicating that EPOR^W439*^ editing did not impair ED (Fig. 2H).

### *Ex vivo* transduction of CD34^+^ cells from healthy donors with integrating HDAd.EPOR^W439*^-LCR-γ-GFP and subsequent transplantation into mice

To assess the engraftment and multilineage differentiation potential of EPOR^W439*^-edited HSPCs, we transduced healthy-donor CD34^+^ cells with HDAd vectors, transplanted them into NCG-X mice, and followed the animals for 16 weeks. Three days before necropsy, mice received clodronate liposomes to deplete mouse macrophages and reduce phagocytosis of human erythroid cells (28,29) (Fig. 3A). EPOR^W439*^ edit installation efficiency, quantified as the combined frequency of the desired TAA and TGA stop-codon alleles, was 1.28% in the infused cells (Fig. S4A), increased to ∼35% in peripheral blood by week 4, and remained stable through week 16 (Fig. 3B). At necropsy, human CD45^+^ chimerism in bone marrow, spleen, and peripheral blood was comparable between mice that received untransduced and EPOR^W439*^-edited cells (Fig. 3C). Similarly, frequencies of CD19^+^ B, CD3^+^ T, and CD14^+^ myeloid cells in bone marrow were comparable between groups, indicating no major lineage skewing following EPOR editing (Fig. 3D).

**Figure 3.**
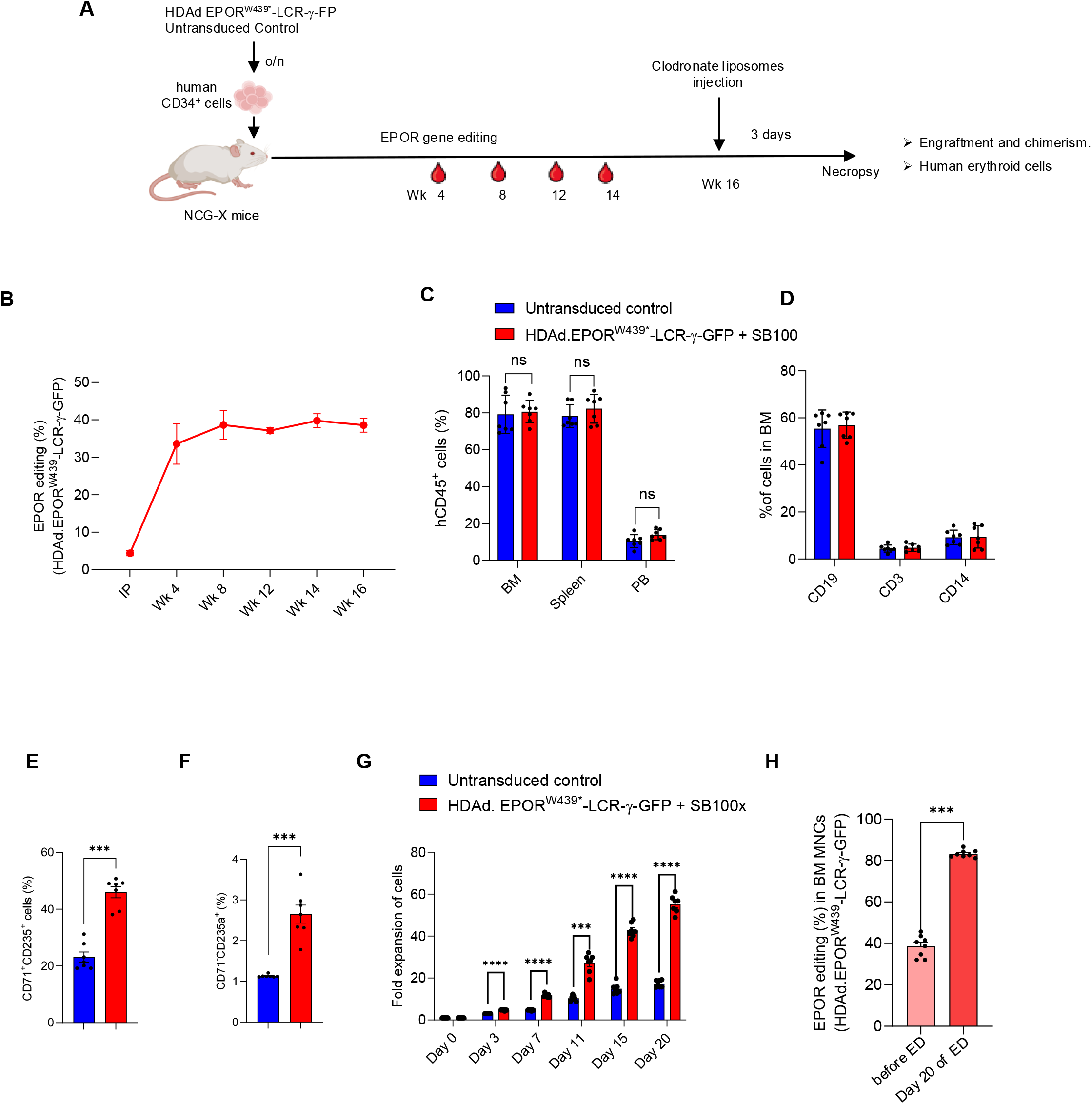
*In vivo* enrichment of EPOR^W439*^edited human hematopoietic cells and enhanced erythroid expansion in humanized mice. **(A)** Experimental design. Human CD34⁺ cells were transduced overnight with HDAd.EPOR^W439*^-LCR-γ-GFP and transplanted into busulfan-conditioned NCG-X mice. Peripheral-blood samples were collected at the indicated time points to monitor EPOR editing. At week 16, mice received clodronate liposomes and were euthanized 3 days later for analysis of human-cell engraftment, hematopoietic chimerism, and erythroid differentiation. Untransduced human CD34⁺ cells served as controls. **(B)** Longitudinal frequency of the EPOR^W439*^ edit in peripheral-blood cells from mice transplanted with HDAd.EPOR^W439*^-LCR-γ-GFP–treated CD34⁺ cells. **(C)** Frequencies of human CD45⁺ cells in the bone marrow (BM), spleen, and peripheral blood (PB) of control and HDAd.EPOR^W439*^-LCR-γ-GFP–treated mice at necropsy. **(D)** Lineage composition of human hematopoietic cells in the BM, determined by the frequencies of CD19⁺ B cells, CD3⁺ T cells, and CD14⁺ myeloid cells. **(E)** Frequencies of human CD71⁺CD235a⁺ erythroid cells in control and HDAd.EPOR^W439*^-LCR-γ-GFP–treated mice. **(F)** Frequencies of human CD71^-^CD235a^+^ erythroid cells in control and HDAd.EPOR^W439*^-LCR-γ-GFP–treated mice. **(G)** *Ex vivo* expansion of BM-derived cells from control and treated mice during culture in erythroid differentiation medium. Fold expansion was calculated relative to the number of cells present at the initiation of culture. **(H)** EPOR^W439*^ editing frequencies in freshly isolated BM cells and after 20 days of erythroid culture, demonstrating preferential expansion of EPOR-edited erythroid cells. Data are presented as mean ± SEM with individual data points shown. Statistical significance was determined using unpaired two-tailed t-test. ns, not significant; **p < 0.01; ***p < 0.001.

Importantly, EPOR^W439*^ edit installation significantly increased erythroid output. The CD71^+^CD235a^+^ early/intermediate erythroid cell population increased from ∼22% to ∼47%, whereas the CD71^−^CD235a^+^ late erythroid cell fraction increased from ∼1% to ∼2.7% (Figs. 3E-F). Amplicon sequencing at necropsy showed EPOR^W439*^ allele frequencies of 31.27% and 37.58% in bone marrow from two mice analyzed (Fig. S4B). Because human erythroid output in this model is limited by factors including restricted marrow space, incompatible mouse erythropoietic growth factors, and phagocytosis of human erythrocytes by mouse macrophages, we subjected bone marrow cells collected at necropsy to *ex vivo* ED. EPOR^W439*^-edited cells expanded ∼55-fold compared with ∼17-fold found for controls by day 20 (Fig. 3G). Consistent with preferential erythroid expansion, EPOR^W439*^ editing enriched from ∼38% in freshly isolated bone marrow cells to >80% after 20 days of ED (Figs. 3H and S4C), demonstrating that HDAd-mediated installation of EPOR^W439*^ in HSPCs preserves engraftment and multilineage differentiation while enhancing erythroid output through preferential expansion of edited cells.

### *In vivo* transduction of humanized mice with integrating HDAd.EPOR^W439*^-LCR-γ-GFP vector

To evaluate EPOR-mediated erythroid enrichment and HbF expression following direct *in vivo* HSPC engineering, we used NSGW41 mice, which provide improved support for human hematopoietic engraftment and erythropoiesis due to the hypomorphic *Kit^W41/W41^* mutation(28). Mice transplanted with healthy-donor human CD34^+^ cells were mobilized and transduced with HDAd.EPOR^W439*^–LCR-γ-GFP + HDAd.SB100x or control HDAd.MGMT/GFP^+^ HDAd.SB100x (Fig. 4A). Mice received clodronate liposomes 3 days before necropsy. Clodronate markedly depleted CD11b^+^Gr-1^low^ macrophage/monocyte populations while largely preserving CD11b^+^Gr-1^+^ cells (Figs. S5A-B).

**Figure 4.**
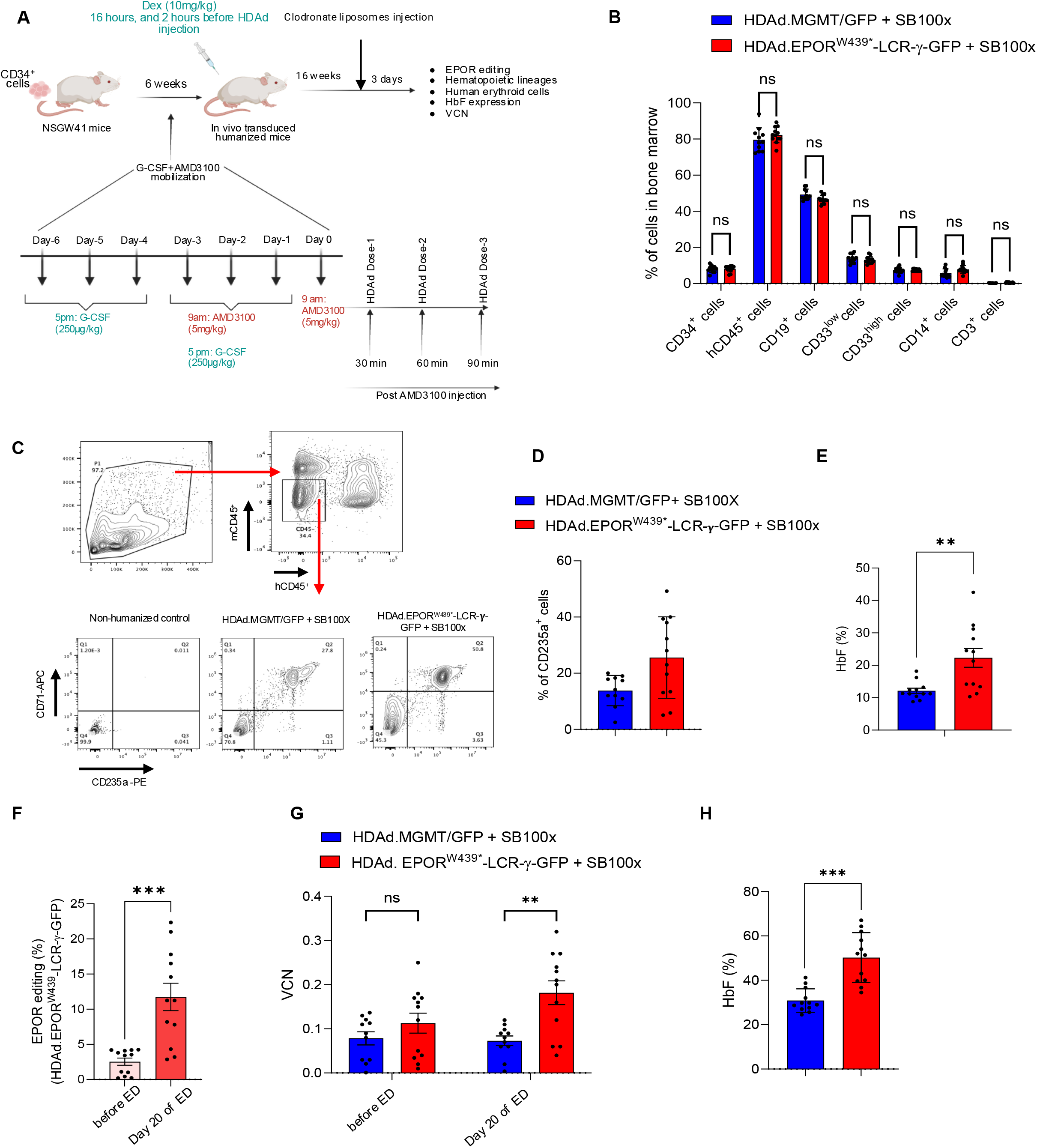
*In vivo* EPOR^W439*^ editing combined with γ-globin gene addition enhances human erythroid output and HbF expression in humanized NSGW41 mice. **(A)** Experimental design. NSGW41 mice were humanized with human CD34⁺cells. Six weeks after transplantation, mice were mobilized with G-CSF and AMD3100 according to the indicated schedule. Dexamethasone (10 mg/kg) was administered 16 hours and 2 hours before HDAd injection. On day 0, mice received three intravenous HDAd doses (8*1010 viral particles in total per animal with 1: 1 ratio of transposase and transposons) at 30, 60, and 90 minutes after the final AMD3100 administration. Sixteen weeks after *in vivo* HSPCs transduction, mice received clodronate liposomes and were analyzed 3 days later for EPOR editing, human hematopoietic-lineage composition, erythroid-cell production, HbF expression, and vector copy number. **(B)** Frequencies of human hematopoietic populations in bone marrow from mice treated with HDAd.MGMT/GFP along with HDAd.SB100x or HDAd.EPOR^W439*^-LCR-γ-GFP along with HDAd.SB100x. Populations analyzed included CD34⁺cells, human CD45⁺ cells, CD19⁺ B cells, CD33^low^ and CD33^high^ myeloid cells, CD14⁺ monocytes, and CD3⁺ T cells. **(C)** Representative flow-cytometric gating strategy used to identify human CD45⁺ cells and CD71/CD235a-defined erythroid populations in bone marrow. Bone marrow from a non-humanized mouse was included as a negative control. Total mouse bone marrow cells were stained with human and mouse CD45 to exclude lymphocytes and human erythroid cells were gated from non-lymphocytes using CD71^+^ and CD235a^+^ cells **(D)** Frequency of human CD235a⁺ erythroid cells in bone marrow of HDAd.MGMT/GFP along with HDAd.SB100x or HDAd.EPOR^W439*^-LCR-γ-GFP along with HDAd.SB100x groups. **(E)** Frequency of HbF-positive cells in freshly isolated bone marrow from mice treated with the HDAd.MGMT/GFP along with HDAd.SB100x or HDAd.EPOR^W439*^-LCR-γ-GFP along with HDAd.SB100x vectors. **(F)** EPOR^W439*^ editing frequencies in freshly isolated bone marrow cells and in erythroid cells generated by ex vivo differentiation of bone marrow cells under regular Epo concentration (3U/ml). **(G)** Vector copy number obtained from total bone marrow cells isolated from bone-marrow and erythroid cells generated from bone marrow cells from the two treatment groups. **(H)** Frequency of HbF-positive cells following *ex vivo* erythroid differentiation of bone-marrow cells. Blue represents HDAd.MGMT/GFP along with HDAd.SB100x, and red represents HDAd.EPOR^W439*^-LCR-γ-GFP plus HDAd.SB100x. Each dot represents an individual mouse; Statistical significance was determined using unpaired two-tailed t-test. ns, not significant; **p < 0.01;***p < 0.001.

At necropsy, 16 weeks after *in vivo* transduction, human CD45^+^ chimerism and CD19^+^, CD33^+^, CD14^+^, and CD3^+^ lineage frequencies were comparable between EPOR^W439*^ and control groups, indicating preserved multilineage hematopoiesis (Fig. 4B; Fig. S5C). Consistent with previous observations, we did not detect human erythroid cells in peripheral blood (Fig. S5D). In addition, peripheral blood hematologic parameters, including hematocrit, were comparable between groups, with no significant differences in leukocyte, erythrocyte, or platelet indices (Fig. S6). In contrast, EPOR^W439*^ treatment increased human erythroid output, with a higher frequency of CD235a^+^ cells in bone marrow (Figs. 4C-D). Importantly, the frequency of HbF-positive cells was significantly increased in the EPOR^W439*^-LCR-γ-GFP group (Fig. 4E), consistent with enrichment of erythroid cells carrying the integrated therapeutic γ-globin cassette.

To further assess preferential expansion of gene-modified erythroid cells, we subjected bone marrow collected at necropsy to *in vitro* ED for 20 days. EPOR^W439*^ edit representation increased from ∼2.5% before differentiation to ∼13% after differentiation (Fig. 4F). This enrichment was accompanied by a significant increase in vector copy number (Fig. 4G), supporting preferential expansion of EPOR^W439*^ cells carrying the SB100x-integrated transgene. Correspondingly, HbF-positive cells increased to ∼50% in the EPOR^W439*–^LCR-γ-GFP group compared with ∼30% in controls (Fig. 4H). Together, these data demonstrate *in vivo* HDAd-mediated EPOR^W439^ edit installation coupled with SB100x-mediated γ-globin integration to preferentially expand gene-modified erythroid cells and increase HbF expression without disrupting multilineage hematopoiesis. Notably, because this mouse model only poorly supports human erythropoiesis (even in the bone marrow), the extent of *in vivo* expansion is limited, and it requires additional *in vitro* ED to reach HbF^+^ cells levels at a therapeutically relevant range.

### *In vitro* transduction studies with CD34^+^ cells obtained from thalassemia and SCD patients

We next evaluated whether EPOR^W439^ edit installation coupled with γ-globin gene addition could preferentially expand therapeutically modified erythroid cells derived from patients with β⁰/β⁰-thalassemia or SCD. In this system, EPOR editing provides an erythroid selective advantage, whereas stable γ-globin expression should correct the functional deficit in disease cells and improve erythropoiesis/survival. Patient CD34^+^ cells were transduced with HDAd.EPOR^W439^–LCR-γ-GFP + HDAd.SB100x or control vectors and, 3 days later, subjected to ED or colony-forming assays (Fig. 5A).

**Figure 5.**
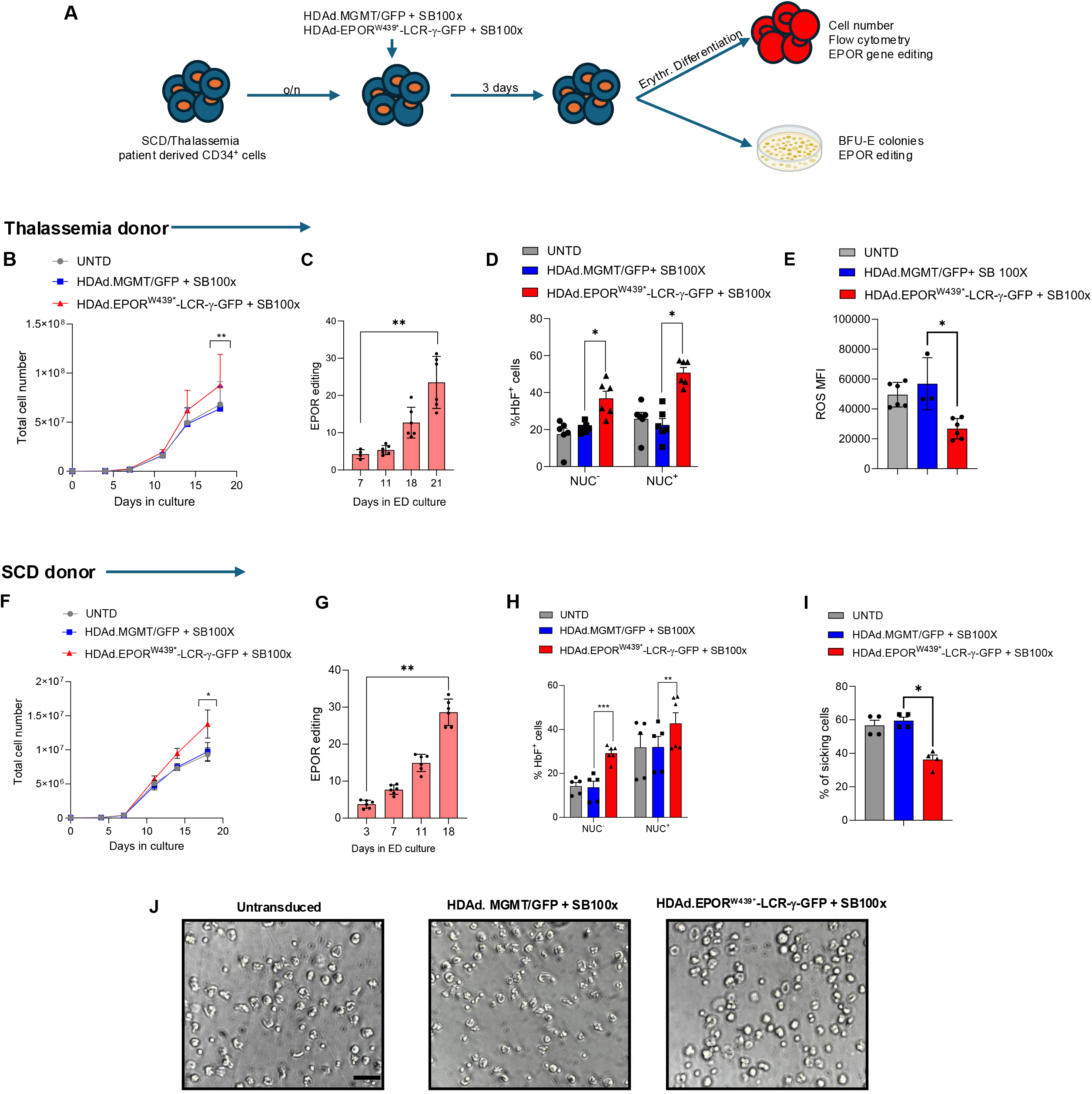
HDAd-mediated EPOR^W439*^ editing combined with SB100x-dependent γ-globin gene addition improves erythroid phenotypes in patient-derived CD34⁺ cells. **(A)** Experimental design. CD34⁺ cells derived from patients with sickle cell disease (SCD) or β-thalassemia were transduced overnight with HDAd.MGMT/GFP plus HDAd.SB100x or HDAd.EPOR^W439*^-LCR-γ-GFP plus HDAd.SB100x (total MOI 4000 vp/cell). Three days after transduction, cells were subjected to erythroid differentiation or plated for colony-forming unit (CFU) assays. Erythroid cultures were analyzed for cell expansion, HbF expression, reactive oxygen species (ROS), and EPOR editing, whereas BFU-E colonies were assessed for EPOR editing. **(B to E)** Studies with β-thalassemia patient-derived CD34⁺ cells. **(B)** Total cell numbers during erythroid differentiation from untransduced control or treated with the indicated vector combinations. **(C)** EPOR^W439*^ editing frequencies at the indicated time points during erythroid differentiation of β-thalassemia cells. **(D)** Frequencies of HbF-positive cells within nucleated (NUC⁺) and enucleated (NUC⁻) erythroid populations derived from β-thalassemia CD34⁺ cells. **(E)** Intracellular ROS levels, reported as mean fluorescence intensity, in β-thalassemia-derived erythroid cells treated with the indicated vector combinations. **(F to J)** Studies with SCD patient-derived CD34⁺ cells. **(F)** Total cell numbers during erythroid differentiation of. **(G)** EPOR^W439*^ editing frequencies at the indicated time points during erythroid differentiation of SCD cells. **(H)** Frequencies of HbF-positive cells within nucleated and enucleated erythroid populations derived from SCD CD34⁺ cells. **(I)** Percentage of sickling erythroid cells generated from SCD patient-derived CD34⁺ cells following the indicated treatments. Gray denotes untransduced cells, blue denotes HDAd.MGMT/GFP plus HDAd.SB100x, and red denotes HDAd.EPOR^W439*^-LCR-γ-GFP plus HDAd.SB100x. **J)** Representative images taken at the end of the sickling assay. The scale bar is 20μm. Statistical significance was determined using unpaired two-tailed t-test. Data are presented as mean ± SEM, with individual measurements shown. *P < 0.05; **P < 0.01.

In β-thalassemic CD34^+^ cells, EPOR^W439*^-treated cells showed greater expansion by the end of ED compared with controls (Fig. 5B). EPOR^W439*^ allele representation progressively increased during differentiation (Fig. 5C), demonstrating preferential enrichment of edited erythroid cells. This enrichment, together with γ-globin gene integration, significantly increased HbF-positive cells in both nucleated and enucleated erythroid populations (Fig. 5D) and significantly reduced intracellular reactive oxygen species (ROS) (Fig. 5E; Fig. S7A). EPOR^W439*^ edit installation also increased BFU-E formation without altering the frequency of CD235a^+^ nucleated erythroid cells (Figs. S7B-C), indicating preserved erythroid maturation. Colony-forming capacity and the frequency of CD235a^+^ nucleated erythroid cells were not adversely affected (Fig. S7D-E). Similarly, EPOR^W439*^ SCD CD34^+^ cells showed increased expansion by the end of ED (Fig. 5F), accompanied by significant enrichment of EPOR^W439*^ allele representation (Fig. 5G). HbF-positive nucleated and enucleated erythroid cells were significantly increased following EPOR^W439*^ edit installation and γ-globin integration (Fig. 5H). Importantly, the proportion of sickled cells was significantly reduced (Figs. 5I-J), demonstrating functional improvement of the SCD phenotype.

### *In vivo* HSC transduction in humanized mice transplanted with HSPCs from a thalassemia patient results in efficient EPOR editing/γ-globin integration and amelioration of disease phenotype

To evaluate *in vivo* therapeutic efficacy in a disease setting, NBSGW mice were humanized with CD34^+^ cells from a patient with β⁰/β⁰-thalassemia (Fig. 6A). Six weeks after transplantation, mice were mobilized and treated with and admixture of HdAd.SB100x and therapeutic HDAd.EPOR^W439*^–LCR-γ-GFP or control HDAd.MGMT/GFP vectors. At necropsy, 16 weeks after *in vivo* transduction, robust human hematopoietic engraftment was detected in bone marrow and spleen. Overall human chimerism and nonerythroid lineage composition were comparable between groups (Fig. 6B-C). Consistent with previous observations, the therapeutic group showed markedly increased human erythroid output, including significantly greater CD235a^+^CD36^+^ and CD235a^+^CD36^−^ populations in bone marrow than the unedited control mice (Fig. 6D). Cytospin-based morphologic analysis further demonstrated more advanced erythroid maturation, with increased late-stage erythroblasts and enucleating cells compared with controls (Fig. 6E). In alignment with preferential enrichment of vector-modified cells, VCN were significantly higher in the therapy group (Fig. 6F). The EPOR^W439^ edit installation efficiency in total bone marrow cells was 19.25 % (Fig. 6G). Importantly, >70% HbF-positive human erythroid cells were found in bone marrow (Fig. 6H), accompanied by reduced spleen- to-body weight ratio and intracellular ROS (Figs. 6I-J) as well as near-complete elimination of splenic hemosiderosis (Fig. 6K), demonstrating substantial amelioration of β-thalassemia-associated phenotypes. The erythroid advantage persisted after explantation, as bone marrow cells harvested from treated animals exhibited greater expansion during 20 days of *ex vivo* erythroid differentiation and generated higher frequencies of human erythroid cells than control bone marrow cells (Figs. S8A-B). Following *ex vivo* ED, EPOR^W439^ allele representation increased further up to 47.5% (Fig. S8C). Colony-forming assays also showed increased BFU-E output (Fig. S8D), while cells pooled from BFU-E contained a markedly higher proportion of HbF-positive cells (∼70% versus ∼15% in controls; Fig. S8E). Clonal analysis using single colonies derived from bone marrow-engrafted cells showed that 19% of colonies had monoallelic and 4% of colonies possessed biallelic EPOR^W439*^ edit installation (Fig. S8F).

**Figure 6.**
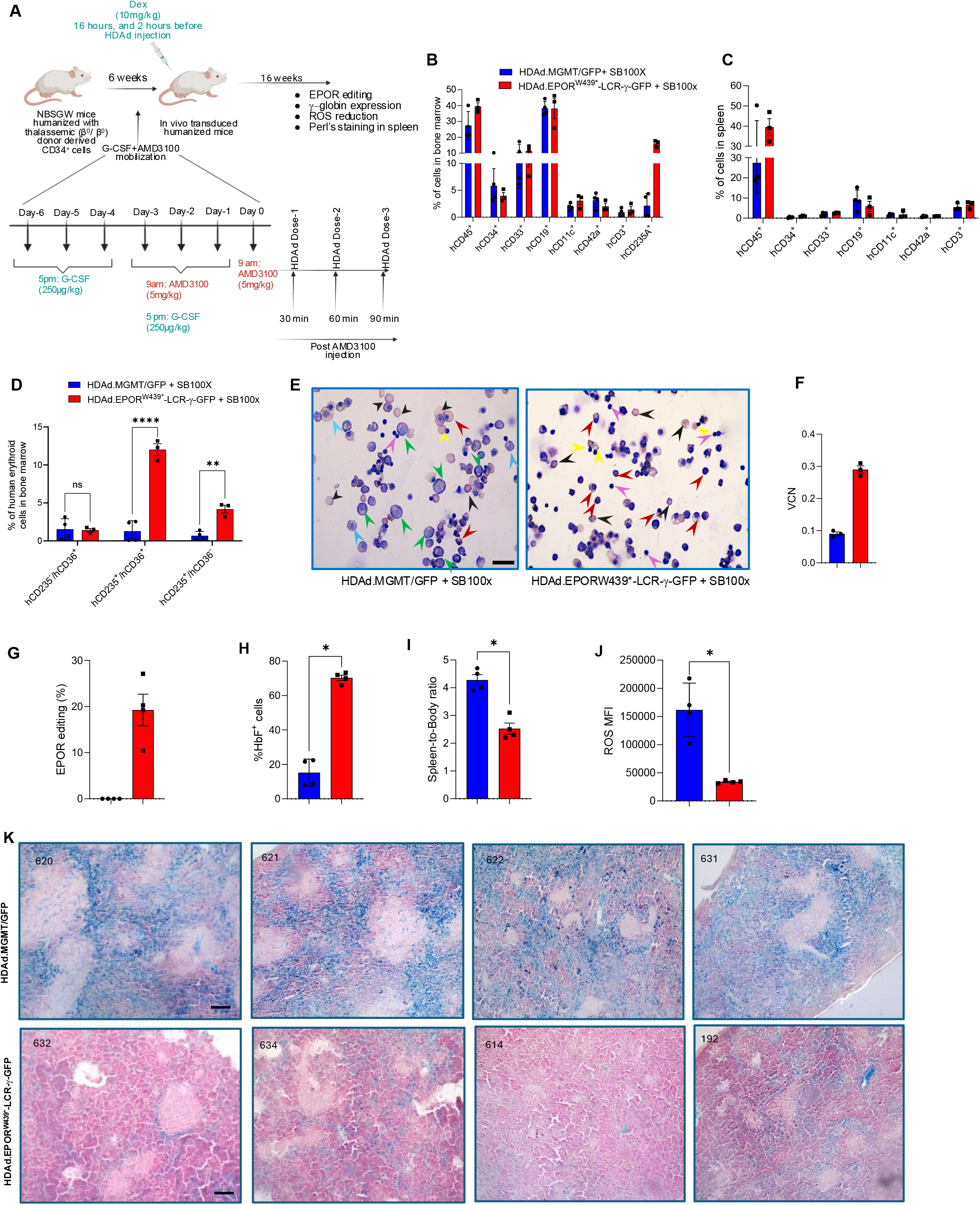
*In vivo* EPOR^W439^ editing coupled with γ-globin gene addition enhances erythroid output and ameliorates disease-associated phenotypes in β-thalassemia patient-derived humanized NBSGW mice. **(A)** Experimental design. NBSGW mice were humanized with CD34^+^ cells from a patient with β⁰/β⁰-thalassemia. Six weeks after transplantation, mice were mobilized with G-CSF and AMD3100 according to the indicated schedule. Dexamethasone (10 mg/kg) was administered 16 and 2 hours before HDAd administration. Following the final AMD3100 injection, mice received three intravenous HDAd doses at 30-minute intervals, for a total dose of 8 × 10^10^ viral particles per animal and a 1:1 ratio of transposon and SB100x vectors. Mice were analyzed 16 weeks after *in vivo* HSPC transduction for hematopoietic engraftment, EPOR editing, erythroid output, γ-globin/HbF expression, oxidative stress, and splenic iron deposition. **(B-C)** Frequencies of the indicated human hematopoietic populations in **(B)** bone marrow and **(C)** spleen of mice treated with HDAd.MGMT/GFP + HDAd.SB100x or HDAd.EPOR^W439*^–LCR-γ-GFP + HDAd.SB100x. **(D)** Frequencies of CD235a/CD36-defined human erythroid populations in bone marrow, demonstrating increased erythroid output in the EPOR^W439*^ treatment group. **(E)** Representative cytospin morphology of bone marrow-derived erythroid cells from the indicated treatment groups. Colored arrows indicate representative erythroid maturation stages. **(F)** Vector copy number (VCN) in total human bone marrow cells, normalized to human RPPH1. **(G)** EPOR^W439*^ editing frequency in total bone marrow cells, determined by targeted next-generation sequencing. **(H)** Percentage of HbF-positive human erythroid cells in bone marrow. **(I)** Spleen-to-body weight ratio at necropsy. **(J)** Intracellular reactive oxygen species (ROS) in bone marrow-derived human erythroid cells, reported as mean fluorescence intensity (MFI). **(K)** Representative Perls’ Prussian blue-stained spleen sections showing iron deposition in control and EPOR^W439*^-treated mice. Blue staining indicates iron accumulation. Data are presented as mean ± SEM, with each symbol representing an individual mouse. Microscopic images shown in panel E and Panel K is taken under 40X magnification. Panels B-D were analyzed by two-way ANOVA with Šídák’s multiple-comparisons test. Panels F-J were analyzed using unpaired two-tailed Student’s *t* tests. ns, not significant; *P* < .05; \**P* < .01; \*\**P* < .001; \**P* < .0001.

Together, these data demonstrate that *in vivo* EPOR^W439^ gene editing coupled with SB100x-mediated γ-globin cassette integration preferentially expands therapeutically modified erythroid cells, increases HbF expression, and ameliorates key disease-associated phenotypes in β-thalassemia humanized mice without disrupting multilineage hematopoiesis.

### Single HDAd vector dual base editing couples EPOR^W439^ selection with HbF reactivation

To eliminate the requirement for SB100x-mediated transgene integration and derive further clinical relevance, we developed a single-vector editing system as a “Hit-and-Run” strategy that simultaneously introduces the EPOR^W439*^ mutation and modulates HbF regulatory sites as a dual base editing system to expand erythroid cells with reactivated HbF. We constructed four HDAd vectors encoding CBE6d and the EPOR^W439*^-targeting sgRNA, each combined with a second sgRNA to *i)* destroy the ATF binding site within the +55-kb enhancer of the *BCL11A* gene, *ii)* replace the -114/-115 BCL11A repressor-binding site within the *HBG1/2* promoters with a TAL1:GATA1 motif recognized by transcriptional activators, *iii)* destroy the LRF binding site (near the -200 bp region) within *HBG1/2* promoters, and *iv)* create a GATA-1 activator binding site in +58kb BCL11a enhancers (Fig. 7A)(30–32).

**Figure 7.**
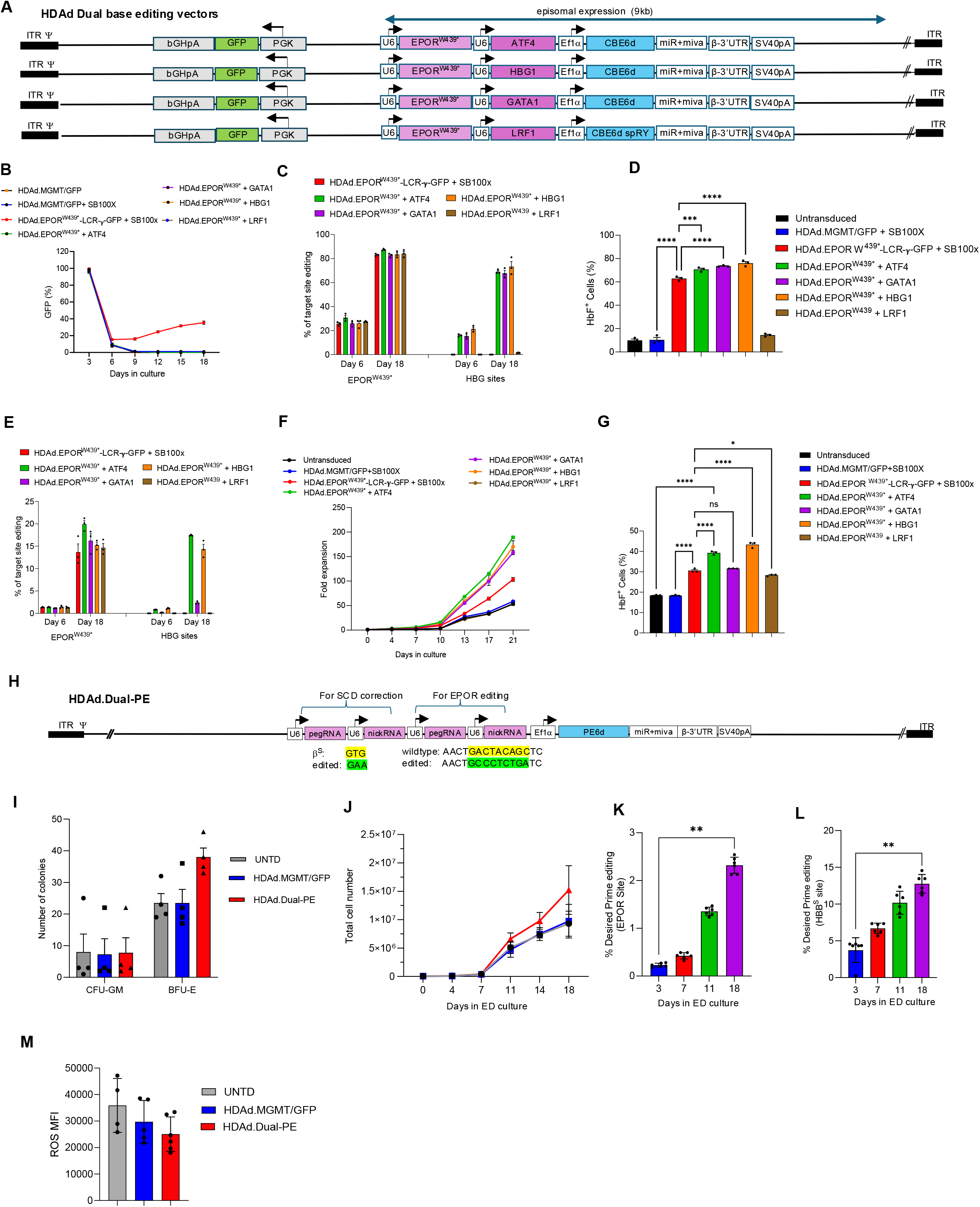
Single-vector dual base and prime editing couples EPOR hypersensitivity with therapeutic editing for hemoglobinopathies. **(A)** Schematic representation of nonintegrating HDAd dual-base-editing vectors. Each vector expresses CBE6d and an EPOR^W439*^-targeting sgRNA together with a second sgRNA targeting the indicated HbF-regulatory element (ATF4, GATA1, HBG1, or LRF1). The editing components are transiently expressed from episomal HDAd genomes. **(B)** Longitudinal GFP marking in HUDEP-2 cells following treatment with the indicated integrating or nonintegrating HDAd vectors. GFP expression declined rapidly with the dual-base-editing vectors, consistent with transient episomal vector expression, whereas marking persisted with the SB100x-integrating control. **(C)** Desired base-editing frequencies at the EPOR^W439*^ site and the corresponding HbF-regulatory target sites on days 6 and 18. **(D)** Percentage of HbF-positive HUDEP-2 cells following treatment with the indicated dual-base-editing vectors. **(E)** Amplicon-sequencing analysis of individual and dual-editing frequencies at EPOR^W439*^ and the corresponding HbF-regulatory sites. **(F)** Fold expansion of healthy-donor CD34^+^-derived cells during erythroid differentiation following treatment with the indicated HDAd vectors. **(G)** Percentage of HbF-positive cells following erythroid differentiation of healthy-donor CD34^+^ cells treated with the indicated dual-base-editing vectors. **(H-L)** Dual prime editing. **(H)** Schematic representation of the HDAd dual-prime-editing vector. The vector expresses PE6d and optimized pegRNA/nicking-sgRNA combinations designed to generate the hypersensitive tEPOR-AL variant and simultaneously correct the sickle mutation in *HBB***. (I)** Numbers of CFU-GM and BFU-E colonies generated from SCD patient-derived CD34+ cells following treatment with the indicated vectors. **(J)** Longitudinal cell expansion during erythroid differentiation of SCD patient-derived CD34+ cells. **(K)** Desired tEPOR-AL prime-editing frequencies during erythroid differentiation. **(L)** Precise correction frequencies of the sickle *HBB* mutation during erythroid differentiation. **(M)** Intracellular reactive oxygen species (ROS) levels following erythroid differentiation, reported as mean fluorescence intensity (MFI). Data are presented as mean ± SEM, with individual symbols representing independent biological replicates. Longitudinal analyses in panels B, C, F, J, K, and L were performed using repeated-measures ANOVA with Šídák’s multiple-comparisons test, as appropriate. Panel I was analyzed by two-way repeated-measures ANOVA with Šídák’s multiple-comparisons test. Endpoint comparisons in panels D, G, and M were performed using one-way ANOVA with Šídák’s multiple-comparisons test. ns, not significant; *P* < .05; \**P* < .01; \*\**P* < .001; \**P* < .0001

In HDAd transduced HUDEP-2 cells, GFP expression rapidly declined following treatment with the nonintegrating dual-editing vectors consistent with loss of episomal HDAd genomes (Fig. 7B). Importantly, EPOR^W439*^ allele representation increased from ∼25%–30% on day 6 to ∼80%–90% by day 18, while editing product representation at HbF-regulatory sites increased from ∼15%–25% to ∼65%–75% (Fig. 7C). Sanger sequencing confirmed efficient edit installation at the GATA1, ATF4, and HBG1 targets, whereas LRF1 editing was undetectable (Figs. S9A-D). ATF4-, GATA1-, and HBG1-targeting vectors generated ∼70%–80% HbF-positive cells, whereas the LRF1-targeting vector failed to induce HbF reactivation due to inefficient editing at the target site (Fig. 7D).

We next evaluated the dual-editing vectors in healthy-donor CD34^+^ cells. Initial transduction efficiencies were comparable among HDAd groups (Fig. S10A), and dual editing did not impair erythroid differentiation, as assessed by CD71 and CD235a expression (Fig. S10B-D). EPOR editing was detected at ∼1.25% on day 6 and increased to ∼15% by day 18, consistent with preferential expansion of EPOR^W439*^-edited cells. Similarly, in cells undergoing combined EPOR^W439*^ and ATF4- or HBG1-targeted editing, the frequencies of ATF4 and HBG1-edited alleles increased from ∼1% on day 6 to ∼14–17% by day 18, consistent with their co-enrichment during EPOR^W439*-^mediated expansion (Fig. 7E). Supporting this selective growth advantage, HSPCs targeted for EPOR^W439*^ editing underwent ∼150- to 190-fold erythroid expansion by day 21 compared with ∼50- to 60-fold expansion of control cells (Fig. 7F). ATF4- and HBG1-targeting vectors generated the highest frequencies of HbF-positive cells, reaching ∼40%–45%. Intermediate or absent response were measured following GATA1 and LRF1 editing due to reduced editing rate (Fig. 7G, Fig. S10E).

Together, these data show that a single nonintegrating HDAd can combine EPOR^W439*^-mediated erythroid expansion with endogenous HbF reactivation.

### Prime editing mediated generation of hypersensitive EPOR

To extend our strategy beyond EPOR^W439*^, we developed a dual-prime-editing HDAd designed to simultaneously correct the sickle mutation in *HBB* and introduce tEPOR-AL, a distinct non-pathogenic truncated EPOR variant first described by Harvey Lodish’s group(33) (Fig. 7H). tEPOR-AL contains a 42-amino-acid C-terminal truncation together with the insertion of two amino acids (AL) immediately before the STOP codon to stabilize the truncated protein. This variant has previously been evaluated using lentiviral vectors in humanized mice and rhesus macaques(34). Unlike EPOR^W439*^, generation of tEPOR-AL requires a precise 9-bp sequence modification. We therefore optimized editing using PE6d by screening two nicking RNAs with three pegRNAs for a 9-bp insertion and two nicking RNAs with five pegRNAs for a 9-bp substitution. The substitution strategy was more efficient, with pegRNA 3 and nicking RNA 2 precisely installing the tEPOR-AL edit with ∼30% efficiency (Figs. S11A-C). Using this combination, we generated an HDAd expressing PE6d, the optimized tEPOR-AL guides, an additional guide pair targeting the sickle *HBB* mutation. In HUDEP-2 cells, the vector precisely installed the tEPOR-AL edit with ∼30% efficiency and resulted in nearly 60% correction of the engineered sickle mutation(35,36), while tEPOR-AL-edited cells progressively enriched during culture, consistent with a selective growth advantage (Figs. S11D-F). We next evaluated the vector in CD34^+^ cells from patients with SCD. Dual prime editing increased BFU-E formation without increasing CFU-GM colonies (Fig. 7I) and enhanced cell expansion during ED (Fig. 7J). Installation of the 9-bp tEPOR-AL modification was less efficient in primary CD34^+^ cells than the single-nucleotide *HBB* correction, highlighting the greater challenge of introducing larger precise edits by prime editing in HSPCs. Nevertheless, by day 18, precise tEPOR-AL edit representation increased to ∼2.3%, whereas correction of the *HBB* sickle mutation reached ∼13% (Figs. 7K-L). Dual prime editing also significantly reduced intracellular ROS (Fig. 7M), supporting functional improvement of the SCD erythroid phenotype. Together, these data demonstrate the feasibility of using a single HDAd prime-editing system to generate a hypersensitive EPOR variant, correct the pathogenic HbS mutation, and preferentially expand therapeutically edited erythroid cells.

### sgRNA-dependent off-target analysis

To evaluate guide-dependent off-target activity, we predicted potential genomic off-target sites for the EPOR^W439*^ base-editing sgRNA and prime-editing guide RNAs using COSMID (https://crispr.bme.gatech.edu/) and evaluated them by targeted amplicon sequencing. For EPOR^W439*^ sgRNA-2, COSMID identified seven candidate off-target loci. C-to-T conversion and InDel frequencies were evaluated at three sites containing cytosines within the relevant base-editing window. We observed approximately 3% conversion at OT-1 in edited samples, compared with ∼2% in the unedited control. At the other two off-target sites, conversion frequencies were similarly low, ranging from 0.1% to 0.3% in both edited and control samples (Figs. S12A-C). For prime editing, we analyzed 11 COSMID-predicted off-target sites for nicking RNA-2, and InDel frequencies remained less than 0.05% across the targets and comparable between control and HDAd.Dual-PE-treated cells (Figs. S13A-B). Similarly, for nine candidate off-target sites for the tEPOR-AL pegRNA spacer, targeted sequencing detected neither the programmed prime-editing product nor increased InDel formation above background (Figs. S14A-C). Together, these results indicate minimal detectable guide-dependent off-target activity at the COSMID-predicted genomic sites examined.

## Discussion

A major challenge for in vivo HSPC gene therapy is achieving therapeutically meaningful levels of gene-corrected cells in diseases where modified HSPCs lack an intrinsic selective advantage. In hemoglobinopathies, our HSPC mobilization– and intravenous HDAd-based platform requires post-treatment enrichment to achieve therapeutically relevant levels of corrected erythrocytes (>20%) (3,4,36–39). Similar chemoselection strategies have been applied to BaEVRless-pseudotyped α-retroviral/lentiviral vectors (40) and CD90-targeted cocal-pseudotyped lentiviral vectors (41). Other delivery platforms, including LNPs, may likewise benefit from an effective *in vivo* enrichment strategy (42).

Here, we established a chemotherapy-free approach that exploits endogenous erythropoietin signaling to preferentially expand therapeutically modified erythroid cells. Recently, Luna et al. showed that CRISPR/Cas9 editing of endogenous *EPOR* to recreate a naturally occurring truncating variant, enhances erythroid output from edited HSPCs during *ex vivo* differentiation(19). Our study extends this concept to *in vivo* HSPC gene therapy by coupling EPOR-mediated erythroid selection with γ-globin gene addition, endogenous HbF reactivation, or correction of the SCD mutation. Although EPOR edit installation efficiency in total bone marrow cells ranged from 4% to 20%, EPOR-edited HSPCs may generate a disproportionately greater erythroid output. Thus, even a relatively small fraction of EPOR-edited HSPCs could potentially give rise to a substantially larger pool of γ-globin-expressing red blood cells, thereby providing meaningful therapeutic benefit.

In β-thalassemia- and SCD-derived CD34^+^ cells, EPOR^W439*^ edit installation combined with γ-globin gene addition produced ∼10-fold enrichment of modified cells during erythropoiesis, increased rates of HbF-positive cells, reduced oxidative stress, and markedly decreased sickling in SCD cultures. Following *in vivo* HSPC transduction, EPOR^W439*^ edit installation increased HbF-positive erythroid cells approximately two-fold in mice humanized with healthy-donor CD34^+^ cells and more than fourfold in β-thalassemia CD34^+^ cell-humanized mice, reaching >70%. Because β-thalassemia is characterized by EPO-driven expansion of erythroid progenitors that inefficiently generate mature erythrocytes(43), we speculate that EPOR^W439*^ edit installation plus γ-globin expression has partially overcome ineffective erythropoiesis, contributing to reduced oxidative stress, splenomegaly/extramedullary erythropoiesis, and splenic iron deposition.

We further developed nonintegrating single-vector strategies combining EPOR-mediated selection with endogenous HbF reactivation by dual base editing or with *HbS* mutation correction by dual prime editing, thereby avoiding SB100x-mediated therapeutic transgene integration. Together, these findings establish EPOR hypersensitivity as a versatile intrinsic selection strategy for in vivo HSPC gene therapy that, when combined with γ-globin expression or SCD-corrective editing, could provide a curative approach for hemoglobinopathies while potentially reducing or eliminating the need for the myeloablative chemotherapy conditioning required for conventional HSCT.

### Limitations of the study

*i)* The long-term consequences of sustained EPOR hypersensitivity remain unknown. Naturally occurring EPOR truncations, including the Mäntyranta variant, indicate that chronic EPOR hypersensitivity can be compatible with a healthy life, but do not establish the safety of engineered EPOR activation in patients with hemoglobinopathies. In β-thalassemia, increased EPO responsiveness plus γ-globin expression may improve ineffective erythropoiesis, whereas in SCD, excessive erythroid expansion could theoretically increase hematocrit and blood viscosity if newly generated erythrocytes are not simultaneously protected from HbS polymerization. Our dual-editing strategies couple EPOR selection with HbF reactivation or *HbS* correction. EPOR-mediated selection may become self-limiting as improved oxygen delivery reduces endogenous EPO levels. Long-term studies should therefore evaluate erythrocyte mass, EPO levels, durability of expansion, and potential thrombotic consequences.
*ii)* In this study, *in vivo* data were generated using EPOR^W439*^ edit installation combined with SB100x-mediated γ-globin integration. Although clinically significant insertional mutagenesis has not been reported with *the Sleeping Beauty transposase* platform, its near-random integration retains theoretical genotoxicity risks. In addition, it requires coadministration of two HDAd vectors and is relatively inefficient in quiescent HSPCs(44–46). These limitations further motivated the development of single-vector, nonintegrating, dual-editing systems. For dual-editing vectors, further optimization is needed to increase simultaneous editing of EPOR and the therapeutic target within the same cell. Under our experimental conditions, robust EPOR-mediated enrichment was observed following biallelic EPOR^W439*^ editing whereas biallelic modification is not necessarily required at the therapeutic loci, as modification of HBG regulatory elements can reactivate HbF and correction of single *Hbb^S^* allele can restore functional β-globin expression(47,48). Prime editing also remains challenging, as the 9-bp tEPOR-AL substitution was substantially less efficient in primary CD34^+^ cells than single-nucleotide *HBB* correction. Although base and prime editors generally reduce DSB formation compared with nuclease editing, unwanted substitutions, InDels, translocations, or other genomic alterations remain possible(49,50), targeted sequencing of COSMID-predicted sites showed minimal detectable guide-dependent off-target activity but cannot exclude unpredicted or guide-independent events(51–53). Further prime editing optimization of the 9-bp tEPOR-AL substitution is also required in the future with e.g., engineered 3’ motifs of epegRNAs, PE8 variants and testing of OptiPrime nominated epegRNA sequences(54–56).
*iii)* Humanized mouse models incompletely reproduce human erythropoiesis. Species-specific differences in the bone marrow microenvironment, cytokines, erythrocyte clearance, and EPO–EPOR signaling limit human erythroid development in xenografts(57–59). This was particularly apparent in healthy-donor CD34^+^ cell-humanized mice, where the strong selective advantage observed under low-EPO conditions *in vitro* became more evident after *ex vivo* ED of recovered bone marrow cells. Studies in human-EPOR transgenic mouse models and nonhuman primates will therefore be important. It is, however, possible that the full potential of our approach will only be realized in models with a hemoglobinopathy background, e.g. hEPOR transgenic thalassemia mice.
*iv)* The use of G-CSF as a mobilizing agent has several disadvantages. G-CSF requires repeated administration, affects the HSC niche, can cause bone pain and splenomegaly, and is contraindicated in SCD(60). We have demonstrated efficient HSPC mobilization and *in vivo* transduction using alternative single-dose regimens, including tGROβ/AMD3100(61) and WU-106/AMD3100,(39) which may be more suitable for future translation.

In summary, EPOR-mediated erythroid selection provides a chemotherapy-free strategy to preferentially expand therapeutically modified erythroid cells, as evidenced by clear improvement of the clinical phenotype in humanized thalassemic mice.

## Acknowledgements

The study was supported by a NIH grant (R01HL130040) (A.L.) and by a grant from the Gates Foundation (INV-017692 (A.L.). Under the grant conditions of the Gates Foundation, a Creative Commons Attribution 4.0 Generic License has already been assigned to the Author Accepted Manuscript version that might arise from this submission. Research reported in this paper was generated using the Department of Laboratory Medicine and Pathology Flow Cytometry Core (CC101467) at the University of Washington.

## Authorship Contributions

A.L. and K.V.K. provided the conceptual framework for the study. A.S.V., K.P., S.G., A.M., C.E.F., R.S., A.L., C.L., H.W., A.K.A. A.G., M.G., E.V., and K.V.K performed the experiments. J.A.Q., N.A., A.A.S., and D.R.L. provided sequences for PE guides. E.Y. provided critical comments on the manuscript. K.V.K. and A.L. wrote the manuscript.

