## Supplementary Methods and Figures for "*In vivo* HSPC gene therapy of hemoglobinopathies without drug selection of corrected cells"

**Reagents:** Prior to transplantation of human $CD34^+$ cells, NSGW41 mice were conditioned with busulfan (Mylan Institutional LLC). For HSPC mobilization and *in vivo* transduction, G-CSF (Neupogen™; Amgen, Thousand Oaks, CA), AMD3100 (MilliporeSigma, Burlington, MA), and dexamethasone sodium phosphate (Fresenius Kabi USA, Lake Zurich, IL) were used.

**Molecular cloning of gene transfer vectors:**

**Lentiviral vectors expressing gene editors and sgRNAs:** A lentiviral plasmid expressing Cas9 (Addgene plasmid #52961) was used to produce Cas9-expressing lentiviral vectors. To generate a lentiviral vector expressing TadCBE, the Cas9 coding sequence was removed by digestion with XbaI and BamHI. The TadCBE coding sequence was PCR-amplified from Addgene plasmid #193835 using primers containing 15-bp overlapping sequences at both ends and inserted into the digested lentiviral backbone using 2× HiFi DNA Assembly Master Mix according to the manufacturer's instructions. To generate lentiviral vectors expressing sgRNAs, Addgene plasmid #57822 was digested with PpuMI and XbaI. Individual sgRNA expression cassettes, including the U6 promoter, were synthesized as gBlocks and ligated into the digested lentiviral backbone using standard T4 DNA ligase–mediated ligation.

Lentiviral vector production: Lenti-X 293T cells were seeded at $9 \times 10^6$ cells per 10-cm dish and allowed to attach overnight. The following day, cells were transfected with 5 µg of transfer plasmid, 2.5 µg of envelope plasmid (Addgene plasmid #12259), and 2.5 µg of gag-pol packaging plasmid (Addgene plasmid #14887). Lentivirus-containing supernatants were collected on day 3 and filtered through a 0.45-µm PES membrane filter. Lentiviral particles were subsequently concentrated using Lenti-X Concentrator (Takara Bio) according to the manufacturer's instructions. The resulting viral pellets were resuspended in Opti-MEM, aliquoted into single-use samples, and stored at −80°C until use.

All base and prime editing sgRNAs used in this study were listed in Supplementary Table 1 and 2.

**Helper-dependent adenovirus vector construction:** HDAd.SB100x, expressing SB100x transposase and FLP recombinase, and HDAd.$MGMT^{P140K}$/GFP were constructed as described previously (1, 2). HDAd.$EPOR^{Q434*}$–LCR-γ-GFP and HDAd.$EPOR^{W439*}$–LCR-γ-GFP were generated through four sequential cloning steps using the pHCA HDAd plasmid backbone. First, the pHCA plasmid (3) was linearized by digestion with PacI and NheI. The optimized EPOR base-editing cassette targeting either Q434* or W439* was PCR-amplified from the corresponding lentiviral plasmid using primers containing 15-bp overlaps with the linearized pHCA backbone and inserted by homology-based DNA assembly. In the second cloning step, the resulting plasmid was linearized with AfeI, and a miR+miva-β-3'UTR-SV40 polyadenylation signal (SV40pA) cassette was PCR-amplified from pBS-EF1α-PEmax-miR+miva-β-3'UTR-SV40pA (4) and inserted into the vector. For the third cloning step, the resulting construct was linearized with PmlI, and a synthetic FRT-IR-bGHpA-GFP-PGK-cHS4 fragment was inserted. This fragment was synthesized as gBlock and contained the indicated FLP recombination target (FRT), *Sleeping Beauty* transposase inverted repeat (IR), bovine growth hormone polyadenylation signal (bGHpA), GFP expression cassette, PGK regulatory element, and cHS4 insulator sequence. Finally, the intermediate plasmid was linearized with AflII, and a fragment containing FRT-IR-µLCR (HS5–HS1)-β-globin promoter-γ-globin regulatory and expression elements was PCR-amplified from HDAd-γ-globin/MGMT (5) and inserted to generate the final pHDAd.$EPOR^{W439*}$–LCR-γ-GFP and pHDAd.$EPOR^{Q434*}$–LCR-γ-GFP plasmids.

To generate HDAd vectors capable of simultaneous EPOR editing and HbF reactivation, a shuttle plasmid containing two sgRNA expression cassettes together with the CBE6d base editor was first constructed. The shuttle construct, designated pBS-U6-EPOR70-2 sgRNA-U6-HbF sgRNA-EF1α-CBE6d-miR+miva-β-3'UTR-SV40pA, was assembled by sequential recombination-based/Gibson cloning into the pBS-Dual plasmid backbone. The U6-EPOR70-2 sgRNA and U6-HbF sgRNA expression cassettes were synthesized as gBlocks. The EF1α promoter and miR+miva-β-3'UTR-SV40pA fragments were PCR-amplified from pBS-EF1α-PEmax-miR+miva-β-3'UTR-SV40pA. The CBE6d coding sequence was PCR-amplified from Addgene plasmid #215822. The four DNA fragments were sequentially assembled into the pBS-Dual vector backbone to generate the complete dual base-editing shuttle plasmid, pBS-U6-EPOR70-2 sgRNA-U6-HbF sgRNA-EF1α-CBE6d-miR+miva-β-3'UTR-SV40pA. The entire dual base-editing cassette was subsequently transferred from the shuttle plasmid into the pHCA HDAd backbone by recombination-based cloning, generating pHCA-U6-EPOR70-2 sgRNA-U6-HbF sgRNA-EF1α-CBE6d-miR+miva-β-3'UTR-SV40pA. The resulting plasmid was then linearized with NheI, and a synthetic PGK-GFP-bGHpA reporter cassette, synthesized as a gBlock, was inserted to generate the final HDAd plasmid, pHDAd-U6-EPOR70-2 sgRNA-U6-HbF sgRNA-EF1α-CBE6d-miR+miva-β-3'UTR-SV40pA-PGK-GFP-bGHpA, as illustrated in Figure 7A.

To construct prime-editing vectors, the PE6d coding sequence was PCR-amplified from Addgene plasmid #207854. The PEmax coding sequence in pBS-EF1α-PEmax-miR+miva-β-3'UTR-SV40pA was replaced with PE6d by restriction digestion and ligation to generate pBS-EF1α-PE6d-miR+miva-β-3'UTR-SV40pA. Individual pegRNA and nickRNA expression cassettes were synthesized as gBlocks and sequentially inserted into the PE6d-containing plasmid to generate U6-pegRNA-U6-nickRNA-pBS-EF1α-PE6d-miR+miva-β-3'UTR-SV40pA constructs. These plasmids were transfected into HEK293 cells to screen different pegRNA/nickRNA combinations and identify the optimal pair for EPOR prime editing. To generate a dual prime-editing construct capable of simultaneously modifying EPOR and correcting the sickle cell disease-causing HBB mutation, the selected EPOR prime-editing construct was further engineered by inserting HBB-specific pegRNA and nickRNA expression cassettes. This generated U6-HBB-pegRNA-U6-HBB-nickRNA-U6-EPOR-pegRNA-U6-EPOR-nickRNA-pBS-EF1α-PE6d-miR+miva-β-3'UTR-SV40pA. To construct the corresponding HDAd dual prime-editing vector, the pHCA backbone was digested with NheI and XmaI to remove an approximately 4.9-kb fragment, and the remaining backbone was re-ligated using T4 DNA ligase. The complete dual prime-editing cassette containing the HBB pegRNA/nickRNA, EPOR pegRNA/nickRNA, EF1α-driven PE6d, miR+miva, β-globin 3'UTR, and SV40 polyadenylation signal was then transferred into the modified pHCA backbone by recombination-based cloning. The resulting construct, designated pHDAd-U6-HBB-pegRNA-U6-HBB-nickRNA-U6-EPOR-pegRNA-U6-EPOR-nickRNA-EF1α-PE6d-miR+miva-β-3'UTR-SV40pA, is shown in Figure 7H.

The integrity and sequence of the completed constructs were confirmed by whole-plasmid nanopore sequencing. Sequence-verified plasmids were subsequently linearized with PmeI for HDAd vector rescue.

HDAd vector rescue and amplification: HDAd5/35++ vectors contained an Ad5-derived capsid in which the Ad5 fiber was replaced with the fiber protein of Ad35. The affinity of the Ad35 fiber for its cellular receptor, CD46, was further enhanced by previously described mutations (6). HDAd6/3+ vectors were derived from adenovirus serotype 6 (Ad6), with the Ad6 fiber knob replaced by the fiber knob of Ad3 (7, 8). The affinity of the Ad3 fiber knob for its receptor, desmoglein 2 (DSG2), was increased by previously described mutations (9).

For HDAd vector production, the corresponding HDAd plasmids were linearized with PmeI and rescued in 116 cells using an Ad6/3+ helper virus. HDAd vectors were subsequently amplified and purified in 116 cells as described previously (10). Residual helper-virus contamination in purified HDAd preparations was <0.05%. Final vector preparations had titers ranging from $3 \times 10^{12}$ to $9 \times 10^{12}$ viral particles (vp)/mL.

**HUDEP-2 cells** were provided by Ryo Kurita (Central Blood Institute, Japanese Red Cross Society, Department of Research and Development, Tokyo, Japan) and Yukio Nakamura (RIKEN BioResource Center, Cell Engineering Division, Ibaraki, Japan). Cells were cultured in medium supplemented with stem cell factor (SCF), erythropoietin (EPO), doxycycline, and dexamethasone, as described previously (11). For HDAd transduction, $0.2 \times 10^6$ cells were seeded in 24-well ultra-low-attachment plates (Corning) at 1 mL of medium per well. HDAd vectors were added at a multiplicity of infection (MOI) of 500–1,000 viral particles (vp) per cell, and the medium was replaced 24 h after transduction. Cells were counted and passaged every 48 h to maintain a density of approximately $0.5 \times 10^6$ cells/mL.

*Lentiviral transduction of HUDEP-2 cells:* HUDEP-2 cells (2 × $10^5$ cells) were resuspended in 1 mL of HUDEP expansion medium supplemented with 8 µg/mL polybrene. Lentiviral vector was added to the cell suspension, followed by spinfection at 800 × g for 30 minutes at room temperature. After transduction, the virus-containing medium was removed, and the cells were washed and reseeded in fresh HUDEP expansion medium. Puromycin selection (2 µg/mL) was initiated 7 days after transduction.

*HUDEP-2 cell culture erythroid differentiation:* Erythroid differentiation (ED) was performed using a previously described protocol with minor modifications. Cells were initially seeded at a density of 1 × $10^5$ cells/mL in IMDM supplemented with GlutaMAX, 3% human AB serum, 2% FBS, insulin (10 µg/mL), heparin (3 U/mL), erythropoietin (EPO; 3 U/mL), holotransferrin (200 µg/mL), stem cell factor (SCF; 100 ng/mL), interleukin-3 (IL-3; 10 ng/mL), and doxycycline (1 µg/mL). On day 3, cells were reseeded at 2 × $10^5$ cells/mL in the same medium. On day 6, cells were reseeded at 4 × $10^5$ cells/mL in differentiation medium containing the same supplements, except doxycycline. On day 8, cells were reseeded at 1 × $10^6$ cells/mL in day 4 medium with the concentration of holotransferrin increased to 500 µg/mL. Cells were collected on day 11 for erythroid and HbF analysis.

*In vitro selection of transduced HUDEP-2 cells:* Cells were treated with 50 µM $O^6$-benzylguanine ($O^6$-BG) for 2.5 hours, followed by 35 µM BCNU (Carmustine) for 1 hour. The drugs were then removed by washing the cells twice with PBS and seeded in fresh medium.)

**$CD34^+$ cells:** Cryopreserved $CD34^+$ cells from mobilized healthy donors were obtained from the Fred Hutchinson Cancer Center (Seattle, WA). $CD34^+$ cells from patients with sickle cell disease (SCD) homozygous for $HBB^{S/S}$ (n = 3) were isolated from exchange-transfusion blood samples collected at George Papanikolaou Hospital (Thessaloniki, Greece). Peripheral blood mononuclear cells (PBMCs) were first purified by Ficoll density-gradient centrifugation, followed by immunomagnetic positive selection of $CD34^+$ cells (Miltenyi Biotec). Approximately 0.5–2.5 × $10^6$ $CD34^+$ cells were recovered from 320–350 mL of steady-state peripheral blood. $CD34^+$ cells from patients with β-thalassemia major ($\beta^0/\beta^0$) were obtained during two mobilization clinical trials conducted at George Papanikolaou Hospital, Thessaloniki, Greece.

*$CD34^+$ cell transduction:* The cells were recovered from frozen stocks and incubated for 24 hours in serum free medium (Stemspan SFEMII, Stemcell Technologies) supplemented with penicillin/streptomycin (Gibco™), Flt3 ligand (Flt3-L, 100 ng/ml), thrombopoietin (TPO, 100 ng/ml) and Stem Cell Factor (SCF, 100 ng/ml) and the small molecules StemRegenin1 (SR1, 1 µM) (Cellagen Technology) and Ly2228820 (Ly, 100 nM) (Selleckchem). All cytokines were obtained from PeproTech. $CD34^+$ cells were transduced with the HDAd5/35-GFP or HDAd6/3-GFP vectors along with the HDAd-SB vector (for integration) in low-attachment plates for 48 hours, at a total MOI 4000 vp/cell, before transferred in erythroid differentiation and methylcellulose-based medium.

*In vitro selection:* HUDEP-2 cells were subjected to chemoselection on day 22 after HDAd transduction. Cells were treated with 50 mM $O^6$BG for 1 hour, followed by 35 mM BCNU for 2.5 hours, for a total treatment duration of 3.5 hours. Drug-containing medium was then removed, and the cells were washed twice with PBS and reseeded in fresh culture medium. Cell viability was monitored every 2 days. During recovery, cells were passaged every 4 days until viability reached 90–100%. Thereafter, cells were maintained at 0.2 × 10^6 cells/mL and passaged every 2 days.

*In vitro erythroid differentiation:* Differentiation of transduced and non-transduced human $CD34^+$ cells into erythroid cells was done based on a 3-step protocol developed by Douay et al. (12) In step 1, cells were cultured at a density of $10^4$ cells/ml for 7 days in a basal medium containing Iscove's modified Dulbecco's medium (IMDM), 5% human plasma, glutamine, Pen-Strep, heparin (2 IU/ml), insulin (10 µg/ml), Holo-Transferrin (330 µg/ml) supplemented with hydrocortisone (1 µM), SCF (100 ng/ml), IL-3 (5 ng/ml) and erythropoietin (EPO) (3 U/ml). In step 2, cells were cultured at a density of $10^5$ cells/ml for 4 days in the same basal medium, as previously, supplemented with SCF (100 ng/ml) and EPO (3 U/ml). Finally, in step 3, the cells were cultured at a density of $10^6$ cells/ml for 7-10 additional days in the same medium, supplemented only with EPO (3 U/ml).

*Colony-forming unit (CFU) cultures:* 2000-3000 human $CD34^+$ cells, or $5x10^4$ human $CD45^+$ cells isolated from chimeric bone marrow were plated in semi-solid methylcellulose-based medium containing cytokines, MethoCult™ H4434 (StemCell Technologies), according to the manufacturer's instructions. After 11-14 days of incubation, CFUs were classified and enumerated under a light microscope by trained operators. Suspensions of pooled colonies were made with at least 25 CFU-GM or BFUE colonies for the analysis of GFP expression.

*HEK293 cell transfection:* Cells were seeded at $1 \times 10^5$ cells per well in 24-well plates and allowed to attach overnight. The following day, the culture medium was replaced with serum-free, antibiotic-free medium. Cells were transfected with 500 ng of plasmid DNA using Lipofectamine LTX at a DNA:Lipofectamine LTX ratio of 1:3, according to the manufacturer's instructions. The transfection mixture was incubated with the cells overnight. Cells were collected on day 4 for analysis of gene-editing efficiency.

**Editing analysis:** Genomic regions encompassing the respective sgRNA target sites were PCR-amplified using locus-specific primers listed in Supplementary Table 3. PCR amplicons were first analyzed by Sanger sequencing to obtain an initial estimate of editing efficiency. Sanger sequencing chromatograms were analyzed using the EditR platform to measure nucleotide substitutions within the sgRNA-targeted region based on changes in chromatogram peak intensities relative to the reference sequence. In some experiments, quantitative analysis, the same genomic regions were subjected to targeted amplicon next-generation sequencing (NGS) Sequencing was performed using illumina MiSeq platform to obtain approximately 50,000–250,000 paired end reads per sample. Raw FastQ sequencing reads were analyzed using CRISPResso/CRISPResso2 with the corresponding reference amplicon sequence and sgRNA sequence for each target, as listed in Supplementary Table 4. Reads were aligned to the reference sequence, and editing frequencies were determined from the nucleotide distribution at the intended target nucleotide(s). For base-editing experiments, the percentage of reads containing the desired nucleotide conversion at the specified target position was used to calculate editing efficiency. For experiments involving multiple targeted nucleotides or bystander edits, nucleotide frequencies at each relevant position were analyzed individually. The completed desired editing was reported as editing efficiency. All FASTQ files have been submitted to the NCBI Sequence Read Archive (SRA) under submission accession SUB16475226 and will be released upon publication.

**Off target site analysis:** *EPOR 70-2 sgRNA:* Potential off-target sites for the EPOR 70-2 sgRNA were identified using COSMID (CRISPR Off-target Sites with Mismatches, Insertions, and Deletions) against the human GRCh38/hg38 genome. The guide-strand sequence, 5′-GTGTCCATGGACGCAAGAGC-3′, was entered in the same orientation as expressed from the U6 promoter, with an NGG PAM specified at the 3′ end. The search included conventional off-target sites containing up to three mismatches, as well as sites containing a single-nucleotide DNA insertion or deletion with up to one additional mismatch. COSMID-designed PCR amplicons were restricted to 250–350 bp, with an optimal length of approximately 300 bp. We curated the output by removing exact duplicate genomic-coordinate entries and collapsing overlapping insertion-, deletion-, or PAM-bulge alignments representing the same genomic locus. For each locus, we retained the representative no-indel alignment with the fewest mismatches and a valid PAM, when available. We analyzed the on-target EPOR locus separately and prioritized the remaining unique candidate off-target loci for targeted amplicon sequencing in matched control and base-edited samples. We isolated genomic DNA from cells treated with the control HDAd.MGMT/GFP vector or the HDAd. $EPOR^{W439*}$-LCR-γ-GFP vector. We amplified each off-target locus by PCR and subjected the resulting amplicons to targeted next-generation sequencing as given in Supplementary Table 5. We analyzed sequencing reads using CRISPResso2 with the corresponding locus-specific amplicon sequence and sgRNA protospacer as references.

*pegRNA-dependent off-target editing analysis:* Potential off-target sites for the EPOR pegRNA spacer (5′-TACCTTGTGGTATCTGACTC-3′) were identified using COSMID against the human GRCh38/hg38 genome assembly. We searched for the spacer with a 3′ NGG PAM, allowing conventional off-target sites with up to three nucleotide mismatches and sites with a single-nucleotide DNA insertion or deletion and up to one additional mismatch. Redundant entries corresponding to overlapping genomic coordinates or alternative mismatch, bulge, or PAM alignments were manually consolidated, resulting in nine unique predicted off-target loci in addition to the on-target EPOR locus.  The oligos and reference sequences were given in Supplementary Table 6

*Nick RNA-dependent off-target editing analysis:* Because the nicking sgRNA does not contain a primer-binding site or reverse-transcription template, off-target activity at these loci was evaluated primarily by quantifying insertions and deletions surrounding the predicted Cas9 nick site rather than by searching for the programmed prime-editing outcome. Indel frequency was calculated as the number of aligned reads containing an insertion or deletion within the CRISPResso2 quantification region divided by the total number of high-quality aligned reads. The oligos and reference sequences were given in Supplementary Table 7. Results were compared between matched control and HDAd.Dual-PE–treated samples from three independent donors. Data are presented as mean ± SEM, with individual donor values shown. We isolated genomic DNA from cells treated with the control HDAd.MGMT/GFP vector or the HDAd. Dual-PE vector. We amplified each off-target locus by PCR and subjected the resulting amplicons to targeted next-generation sequencing. We analyzed sequencing reads using CRISPResso2 with the corresponding locus-specific amplicon sequence and sgRNA protospacer as references. Editing frequencies were compared between matched HDAd.MGMT/GFP and HDAd.Dual-PE samples from three independent donors. Data are presented as mean ± SEM, with individual points representing independent donors.

**VCN:** Vector copy number (VCN) was determined by droplet digital PCR (ddPCR) using genomic DNA isolated from mouse samples. Genomic DNA was extracted using the PureLink Genomic DNA Mini Kit and quantified with the Qubit dsDNA High Sensitivity Assay Kit (Cat. No. Q33231; Thermo Fisher Scientific). Fifty nanograms of genomic DNA were used as template per ddPCR reaction. An EGFP-specific primer/probe set was used to quantify vector copies, and a human RPP30-specific primer/probe set was used as the endogenous reference as given in Supplementary Table 7.

Reactions were prepared using ddPCR Supermix for Probes (No dUTP) (Cat. No. 186-3023; Bio-Rad). Droplets were generated with a QX200™ Droplet Generator, followed by PCR amplification according to standard ddPCR procedures. Amplified droplets were analyzed using a QX200™ Droplet Reader, and absolute target concentrations were calculated using the associated Bio-Rad analysis software. VCN was calculated using the following formula: VCN per $GFP^+$ cell = bulk-population VCN / fraction of $GFP^+$ cells. This calculation assumes two copies of RPP30 per diploid mouse genome. Genomic DNA from nontransduced mice and no-template controls were included in each experiment.

For bone marrow analyses, genomic DNA was isolated from total nucleated bone marrow cells. Accordingly, the reported VCN represents the average number of vector copies per diploid genome in the bulk bone marrow cell population and was not corrected for the percentage of $GFP^+$ cells unless otherwise indicated. When flow-cytometric data from the corresponding sample were available, the estimated VCN per $GFP^+$ cell was calculated by dividing the bulk-population VCN by the fraction of $GFP^+$ cells. This value was reported as an estimate because the calculation assumes that vector copies are confined to $GFP^+$ cells, whereas GFP-negative cells may contain transcriptionally inactive, incomplete, or transposase-independent vector sequences. Therefore, bulk VCN and GFP marking were presented separately, and corrected values were explicitly identified as estimated rather than directly measured VCN per $GFP^+$ cell.

**Mice:** All experiments involving animals were conducted in accordance with the institutional guidelines set forth by the University of Washington. The University of Washington receives accreditation from the Association for the Assessment and Accreditation of Laboratory Animal Care International (AALAC) and all live animal work conducted at the university is in accordance with the Office of Laboratory Animal Welfare (OLAW) Public Health Assurance (PHS) policy, USDA Animal Welfare Act and Regulations, the Guide for the Care and Use of Laboratory Animals and the University of Washington's Institutional Animal Care and Use Committee (IACUC) policies. The studies were approved by the University of Washington IACUC (Protocol No. 3108-01).

***Ex vivo* HSPC transduction and transplantation into NCG-X mice:** Adult human $CD34^+$ cells were transduced overnight with HDAd.$EPOR^{W439*}$–LCR-γ-GFP at a multiplicity of infection (MOI) of 2,000 viral particles per cell. The following day, cells were collected, washed with PBS, and resuspended at $1 \times 10^6$ cells in 100 µL PBS per mouse. Cells were transplanted by tail-vein injection into busulfan-conditioned NCG-X mice. The busulfan dose was 12.5mg/kg.

Peripheral blood was collected at the indicated time points after transplantation to monitor the persistence of EPOR-edited human cells. At 16 weeks post-transplantation, mice were euthanized, and bone marrow, peripheral blood, and splenic cells were harvested for analysis of human hematopoietic engraftment. Human and mouse hematopoietic cells were distinguished by flow cytometry using antibodies against human CD45 (hCD45) and mouse CD45.1 (mCD45.1). Human-cell engraftment was calculated as: Human engraftment (%) = [hCD45 / (hCD45 + mCD45.1)] × 100. To determine EPOR editing in engrafted human cells, genomic DNA was extracted using QuickExtract DNA Extraction Solution (Lucigen; Cat. No. QE0905T). The human EPOR target region was amplified using human-specific EPOR primers and subjected to Sanger sequencing followed by EditR analysis to quantify editing efficiency.

*Macrophage depletion with clodronate liposomes:* Macrophages were depleted by intravenous administration of 200 µL clodronate-containing liposomes (5 mg/mL; ClodronateLiposomes.com) per mouse. Control animals received an equivalent volume (200 µL) of PBS or control liposomes. Clodronate liposomes were administered intravenously at the indicated time point before subsequent experimental procedures. To confirm macrophage depletion, peripheral blood was collected 72 hours after clodronate-liposome administration and analyzed by flow cytometry using macrophage/monocyte-specific surface markers. The frequency of circulating macrophage/monocyte populations after treatment was compared with the corresponding pretreatment baseline for each animal to determine the extent of depletion.

**Mobilization and *in vivo* transduction of human $CD34^+$ cells in a humanized NSGW41 mouse model:** The immunodeficient NOD.Cg-Kit W-41J Prkdc scid Il2rg tm1Wjl /WaskJ (NSGW41) mice were generously provided by Thalia Papayannopoulou (University of Washington). A humanized model was generated by transplanting human $CD34^+$ cells from healthy donors into NSGW41 mice ($1x10^6$/recipient), post partial myeloablation (Busulfan 12.5 mg/kg). Six weeks post transplantation, the mice, having a human bone marrow chimerism, were mobilized by a 7-day mobilization scheme, including G-CSF 250 µg/kg s.c. (days 1-6) and AMD3100 5 mg/kg i.p. (days 5-7), as previously described (13). Forty minutes after the last AMD3100 injection, mice received i.v. the HDAd vectors at a total dose of $8x10^{10}$ viral particles (divided into two doses, 30 min apart). Sixteen and two hours before i.v. injection of HDAd vectors, the animals received Dexamethasone (i.p, 10 mg/kg). One month post *in vivo* transduction, the animals were injected i.p. with freshly prepared $O^6BG$ (30 mg/kg, in two doses, 30 min apart) and 5mg/kg BCNU for the *in vivo* selection of transduced cells. One month post *in vivo* selection, NSGW41 mice were sacrificed, and their hematopoietic tissues were collected, for assessment of multilineage engraftment and transduction efficiency. Human $CD45^+$ cells were isolated from the bone marrow and transplanted into secondary NSGW41 recipients. A small fraction of these cells was used for CFU cultures.

**Xenotransplantation and in vivo transduction studies with patient $CD34^+$ cells:** For the *in vivo* studies, 1 × $10^6$ $CD34^+$ cells obtained from a $\beta^0/\beta^0$-thalassemic patient were transplanted into NBSGW mice. Human hematopoietic cells engraftment was confirmed 6 weeks after transplantation by fluorescence-activated cell sorting (FACS). Following successful engraftment, mice were mobilized using a combination of granulocyte colony-stimulating factor (G-CSF) (for 6 days) and plerixafor (for 4 days) according as described above. Sixteen and two hours before *in vivo* transduction 10mg/kg Dexamethasone was administrated. Upon completion of the mobilization regimen, animals received an intravenous injection of the HDAd6/3+ $EPOR^{W349*}$-LCR-γ-GFP vector + HDAd6/3+ SB100X or HDAd6/3+ MGMT/GFP + HDAd6/3+ SB100X at a total dose of $8 \times 10^{10}$ viral particles. Mice were sacrificed 16 weeks after $CD34^+$ cell transplantation, and spleens were harvested for size assessment and histology analysis. Bone marrow samples were subsequently analyzed to determine the extent of human engraftment, multilineage hematopoietic reconstitution, and fetal hemoglobin (HbF) expression in engrafted human erythroid cells. A portion of chimeric bone marrow cells were seeded in erythroid differentiation medium as well as in methylcellulose medium.

**Flow cytometry analyses:** To evaluate the HSC phenotype post transduction, cells were washed and stained with the following fluorochrome-conjugated antibodies against human antigens: CD34-APC, CD38-PE, CD90-PerCP and CD45RA-APC-H7 (BD Biosciences, San Jose View, CA). For the follow up of human $CD34^+$ cell differentiation into erythroid or myeloid cells the following antibodies were used: For erythroid cells: CD235a-PE (BD Biosciences) and CD71 FITC (BD Biosciences); for myeloid cells: CD33-PE (BD Biosciences). To assess the multilineage engraftment of human $CD34^+$ cells in the bone marrow of NSGW41 mice post transplantation, the following antibodies were used: CD45-APC (BD Biosciences), CD19-PerCP (BioLegend), CD3-FITC (BioLegend), CD33-PE (BD Biosciences) and CD235a-PE (BD Biosciences). To measure the percentage of human HSPCs in peripheral blood of NSGW41 mice post mobilization, the following antibodies were used: CD45-APC (BD Biosciences, San Jose View, CA), CD34-PE (BioLegend) and CD90-PerCP (BD Biosciences, San Jos bvyie, CA). Blood collected retro-orbitally at various time points post mobilization and stained with the antibodies for 15 minutes in cool and dark place. After the incubation, erythroid cells were lysed using an RBC Lysis Buffer (BioLegend), followed by a washing step in FACs buffer. After wash, cells were resuspended in FACS buffer and analyzed using a BD FACSymphony™ A3 Cell Analyzer (BD Biosciences, San Jose, CA). Debris was excluded using a forward scatter-area and sideward scatter-area gate. Flow cytometry data were then analyzed using FlowJo (version10.0.8, FlowJo, LLC). A complete list of antibodies is provided in Supplementary Table 9.

**Intracellular flow-cytometric analysis of HbF expression:** Approximately 1 × $10^6$ cells were collected and fixed in 4% formaldehyde in PBS for 15 min at room temperature. Following centrifugation, cells were sequentially permeabilized on ice with 1:1 acetone: water for 3 min, 100% acetone for 3 min, and 1:1 acetone:water for an additional 3 min, with centrifugation between each step. Cells were subsequently resuspended in PBS containing 0.5% BSA.  Permeabilized cells were stained with a PE-conjugated anti-HbF antibody in 100 µL staining buffer for 15–20 min at 4°C, protected from light. Cells were then washed with PBS containing 0.5% BSA and resuspended for flow-cytometric analysis. Where indicated, NucRed Live 647 ReadyProbes Reagent (Invitrogen, R37106) was used to distinguish nucleated from enucleated erythroid cells. Unstained samples and appropriate HbF-negative erythroid cells were used to establish gating.

**Cytospin preparation:** During erythroid differentiation, 0.3–0.5 × $10^5$ transduced and untransduced β-thalassemic and SCD cells were collected for morphological analysis. Cytospins were prepared by centrifugation at 100 × g for 5 min using a ROTOFIX 32 cytocentrifuge (Hettich Zentrifugen). After air drying, slides were stained with May–Grünwald for 5 min followed by Giemsa (Merck, Darmstadt, Germany) for 15 min and subsequently examined microscopically.

**Analysis of Reactive Oxygen Species (ROS) levels:** Intracellular ROS production was assessed in *ex vivo*–differentiated β-thalassemic erythroid cells using the CellROX™ Deep Red Flow Cytometry Assay Kit (Invitrogen, Thermo Fisher Scientific). A total of 1 × $10^6$ cells were incubated with 500 nM CellROX™ reagent for 30 min at 37°C, protected from light. ROS levels were determined by measuring the oxidation-dependent increase in APC fluorescence using flow cytometry.

**Sickling assay:** For sickling assays, erythroid cells were adjusted to a density of 5 × $10^5$ cells/500 µL in stage III erythroid differentiation medium. Sickling was induced by adding an equal volume of freshly prepared 2% sodium metabisulfite in PBS, followed by incubation at room temperature for 30 min.

**Perls staining:** Perls' Prussian Blue staining was carried out using the Perl stain Kit (Atom Scientific). Following deparaffinization and rehydration, tissue sections were incubated in potassium hexacyanoferrate for 5 min and then exposed to a potassium hexacyanoferrate/hydrochloric acid solution for 1 min. After washing in distilled water, slides were mounted. Images were taken using Nikon microscope under 40X magnification as mentioned figure legends.

**Blood analysis:** Blood samples were collected into EDTA-coated tubes and analysis was performed on a HemaVet 950FS (Drew Scientific, Waterbury, CT).

**Statistics.** Data are presented as mean ± SEM. Details of the statistical analyses are provided in the corresponding figure legends. All statistical analyses were performed using GraphPad Prism version 10.0.3 (GraphPad Software, Boston, MA, USA). *P* values < .05 were considered statistically significant.

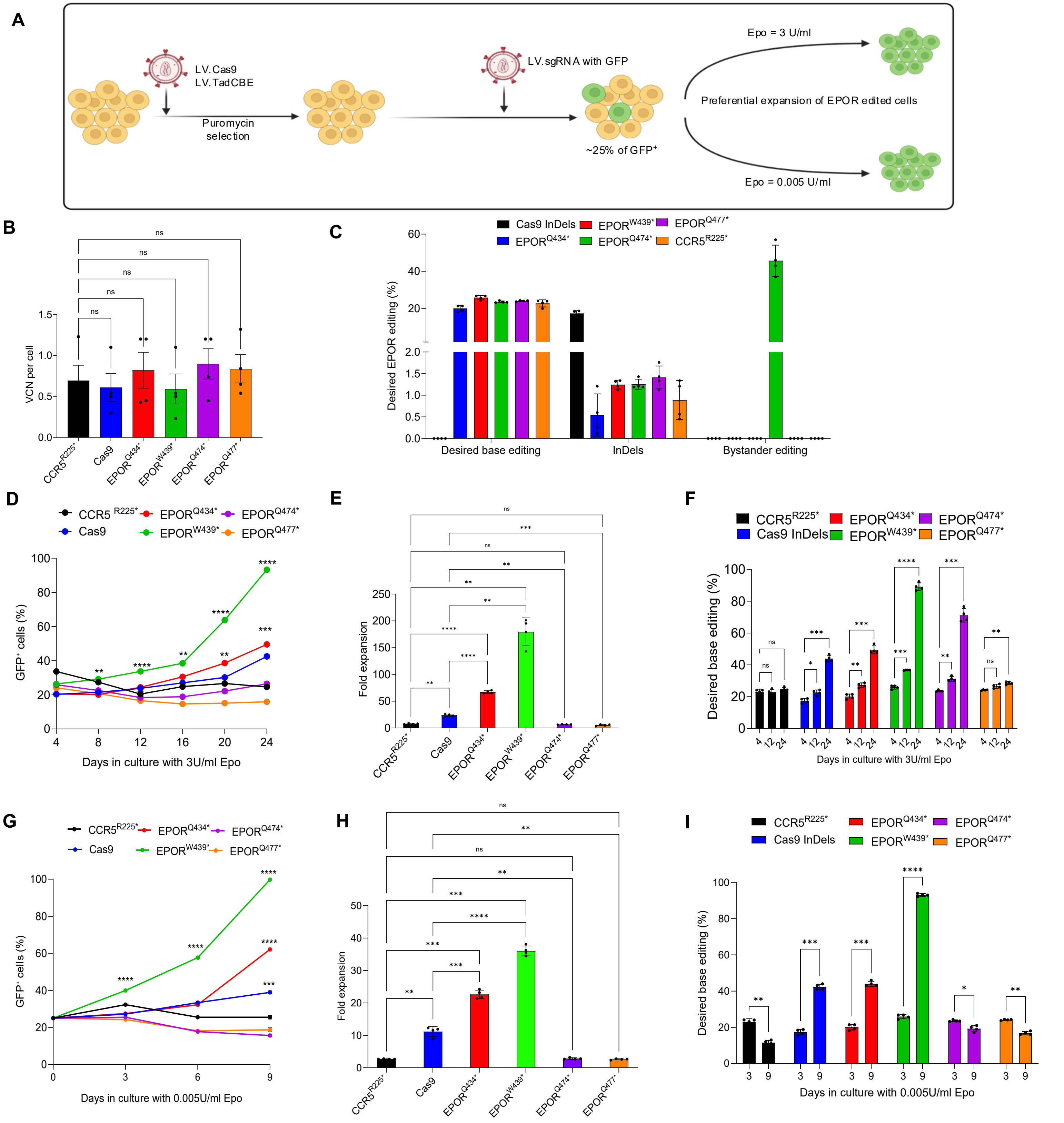


**Supplemental Figure 1. Screening of hypersensitive EPOR variants in HUDEP-2 cells by lentivirus vector-mediated base editing. (A)** Experimental workflow. HUDEP-2 cells expressing Cas9 or the TadCBE adenine base editor were generated and selected with puromycin. Cells were subsequently transduced with lentiviral vectors expressing GFP together with the indicated sgRNAs, resulting in an initial GFP-positive fraction of approximately 25%. The cells were then cultured in medium containing either 3 or 0.005 U/mL EPO to assess the selective expansion of EPOR-edited cells under standard and reduced EPO conditions. **(B)** Vector copy number per cell in cells transduced with sgRNAs targeting $CCR5^{R225}$, $EPOR^{Q434}$, $EPOR^{W439}$, $EPOR^{Q474}$, or $EPOR^{Q477}$, or with the Cas9-indel control. **(C)** Frequencies of the intended base-editing products, indels, and bystander edits generated by the indicated sgRNAs. The discontinuous y-axis permits visualization of both high-frequency intended edits and lower-frequency indel and bystander-editing events. **(D)** Longitudinal analysis of GFP-positive cells cultured in medium containing 3 U/mL EPO.. **(E)** Fold expansion of the indicated cell populations after 24 days of culture in medium containing 3 U/mL EPO. **(F)** Frequencies of the intended base-editing products in the indicated cell populations on days 4, 12, and 24 of culture in medium containing 3 U/mL EPO. **(G)** Longitudinal analysis of GFP-positive cells cultured under reduced-EPO conditions containing 0.005 U/mL EPO. **(H)** Fold expansion of the indicated cell populations after 9 days of culture in medium containing 0.005 U/mL EPO. **(I)** Frequencies of the intended base-editing products in the indicated cell populations on days 3 and 9 of culture in medium containing 0.005 U/mL EPO. **(I)** Frequencies of the desired EPOR edits or Cas9-induced indels at the indicated time points during culture in 0.005 U/mL EPO. $CCR5^{R225*}$ served as a non-EPOR-targeting base-editing control. Data are shown as mean ± SEM. Statistical significance was determined by Two-Way ANOVA with Dunnet multiple comparisons. ns indicates not significant; *P < .05, **P < .01, ***P < .001, ****P < .0001

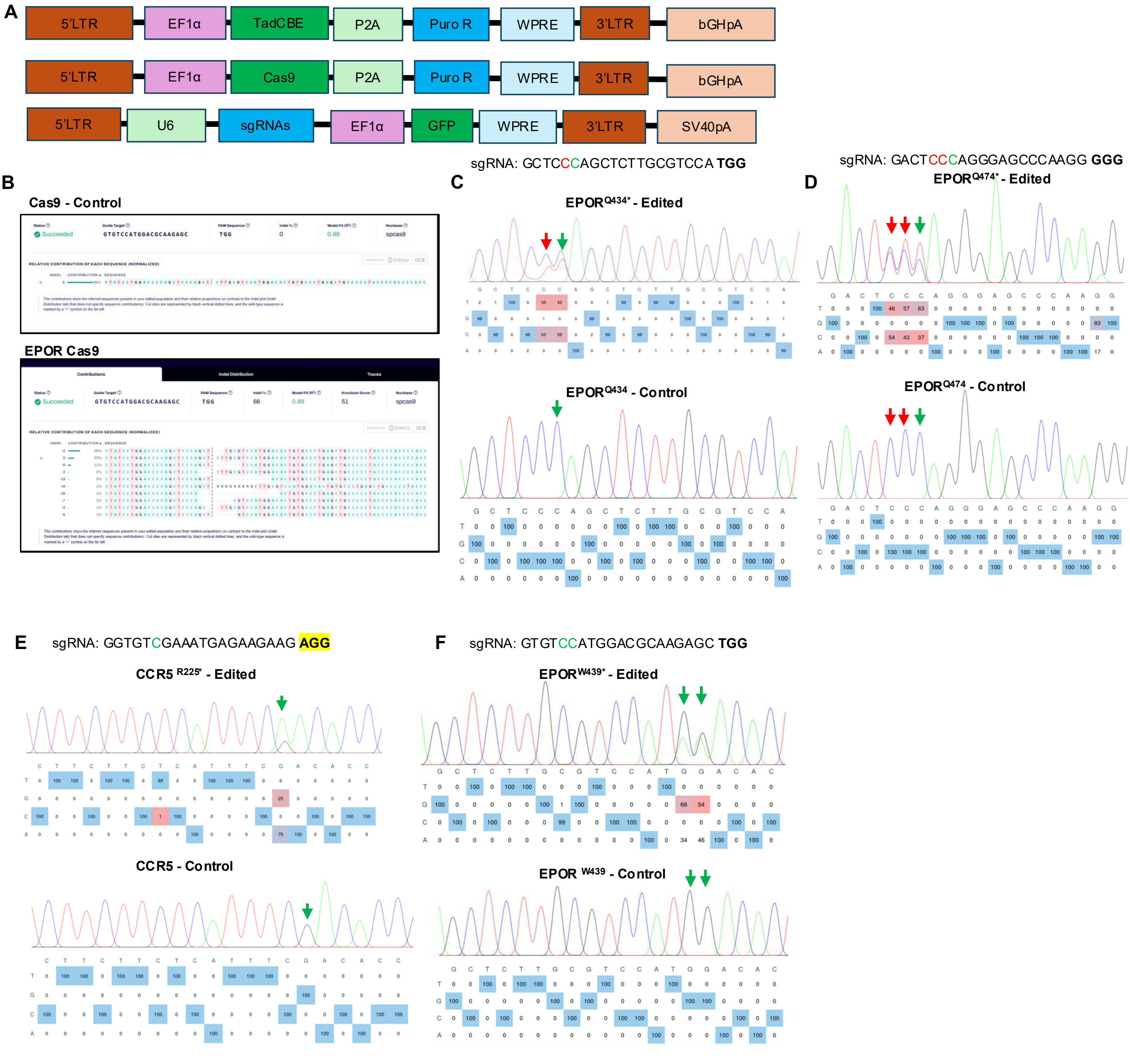


**Supplemental Figure 2. Lentiviral vector design and sequence validation of EPOR- and CCR5-targeted genome editing in HUDEP-2 cells. (A)** Schematic representation of lentivirus constructs used for stable expression of genome editors. The constructs include an EF1α-driven TadCBE or Cas9 linked to a puromycin resistance cassette (P2A-PuroR), flanked by 5′ and 3′ LTRs, WPRE, and polyadenylation signals (Top panel). Schematic of sgRNA expression cassette. sgRNAs targeting EPOR or control loci are expressed under the U6 promoter, with EF1α-driven GFP enabling tracking of transduced cells (Bottom). **(B)** Representative sequence-decomposition analysis of the EPOR target site in control and Cas9/EPOR-sgRNA–treated cells, demonstrating Cas9-mediated insertion and deletion formation. **(C-F)** Representative Sanger-sequencing chromatograms and nucleotide-frequency analyses of cells edited with sgRNAs targeting EPOR$^{Q434*}$ (C; sgRNA, 5′-GCTCCCAGCTCTTGCGTCCA-3′; PAM, TGG), EPOR$^{Q474*}$ (D; sgRNA, 5′-GACTCCCAGGGAGCCCAAGG-3′; PAM, GGG), CCR5$^{R225*}$ (E; sgRNA, 5′-GGTGTCGAATTGAGAAGAAG-3′; PAM, AGG), EPOR$^{W439*}$ and (F; sgRNA, 5′-GTGTCCATGGACGCAAGAGC-3′; PAM, TGG), together with the corresponding unedited controls. Green arrows indicate the intended nucleotide substitutions, whereas red arrows indicate bystander edits within the editing window. Nucleotide-frequency tables below each chromatogram show the relative frequencies of the four nucleotides at each analyzed position.

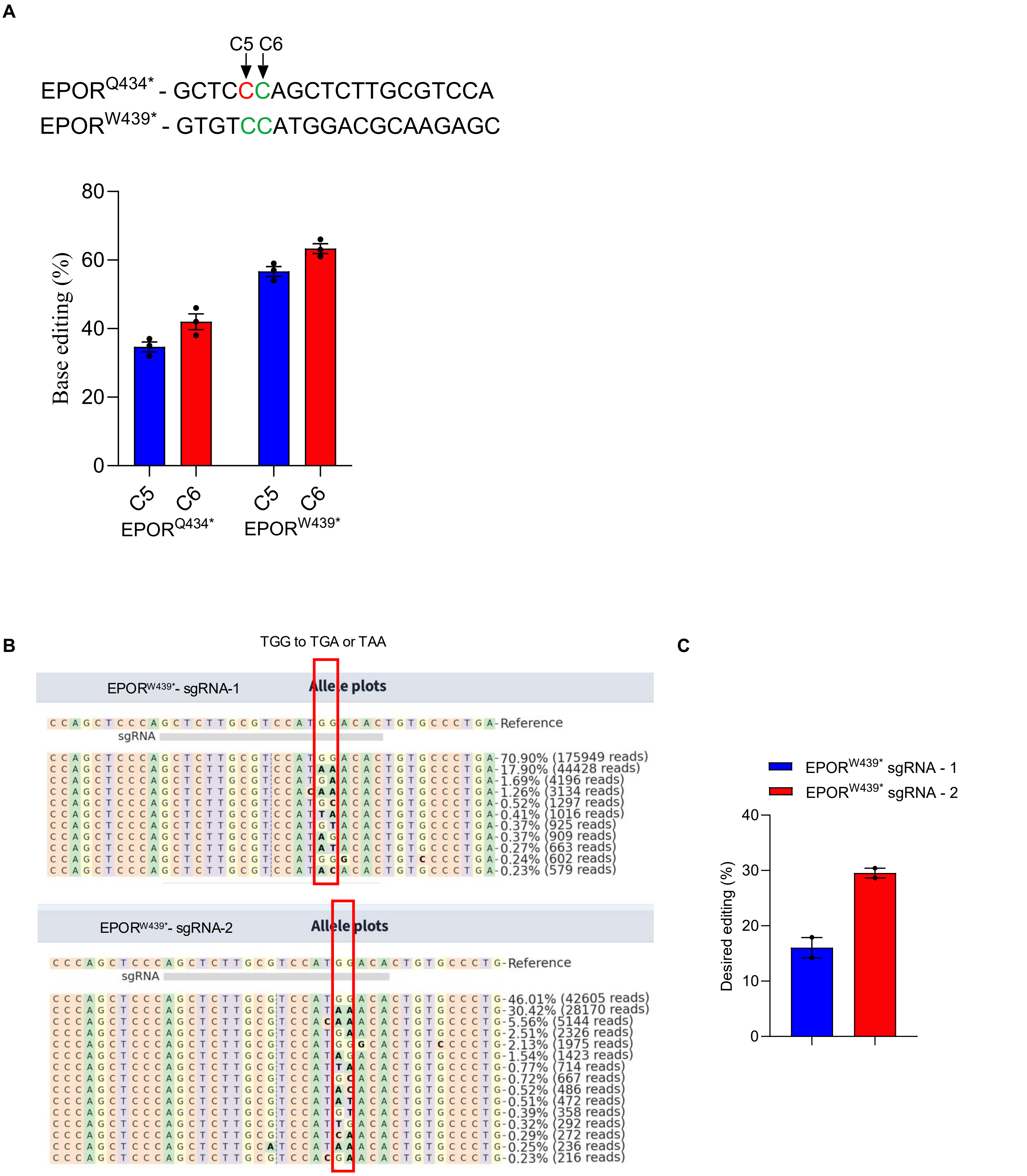


**Supplemental Figure 3. Comparison of two sgRNAs for base editing of the EPOR$^{W439*}$ site.** (A) Target sequences and cytosine base-editing efficiencies at the EPOR$^{Q434*}$ and EPOR$^{W439*}$ sites. Editing frequencies at cytosines C5 and C6 are shown. Among the two variants, the EPOR$^{W439*}$ sgRNA was selected because it achieved the desired target edit without the bystander editing observed with the EPOR$^{Q434*}$ sgRNA **(B)** Representative allele plots from targeted amplicon sequencing of HUDEP-2 cells edited using EPOR$^{W439*}$ sgRNA-1 or sgRNA-2. The reference sequence is shown above each plot, with the sgRNA-binding region indicated. The most abundant edited alleles and their corresponding read frequencies are displayed; nucleotide substitutions relative to the reference sequence are highlighted. **(C)** Quantification of the intended EPOR$^{W439*}$ editing frequency obtained with sgRNA-1 and sgRNA-2. Editing efficiency was calculated as the percentage of sequencing reads containing the desired EPOR$^{W439*}$ nucleotide substitution. EPOR$^{W439*}$ sgRNA-2 produced higher desired editing than sgRNA-1. Bars represent the mean with error bars, and individual replicate values are shown.

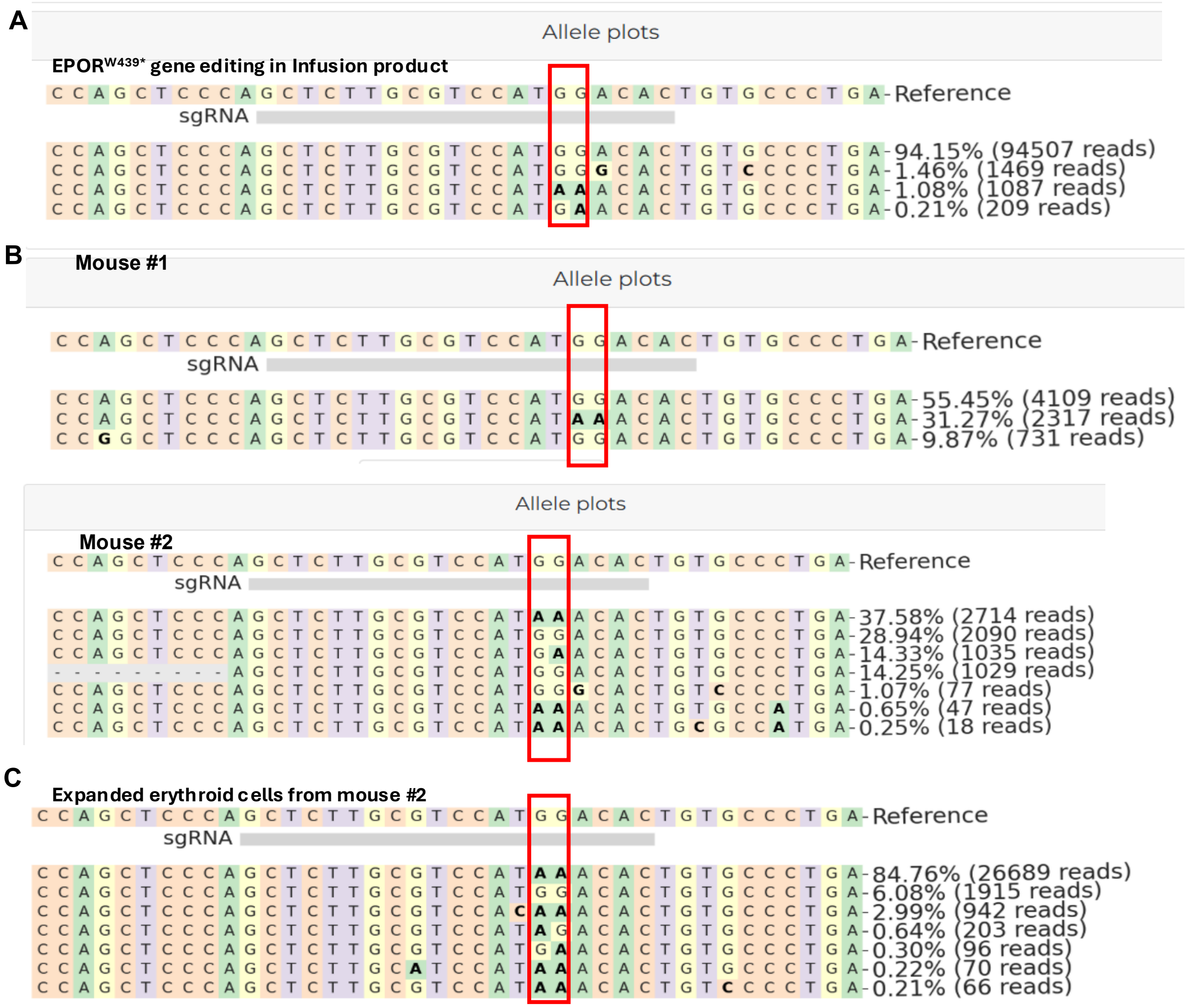


**Supplemental Figure 4.** ***In vivo*** **enrichment of tEPOR-edited alleles following transplantation and erythroid differentiation. (A)** Representative allele plots from targeted amplicon sequencing showing the EPOR target region in the human CD34$^+$ cell infusion product. **(B)** Representative allele plots of bone marrow cells recovered from mice #1 and #2 at 16 weeks after transplantation. **(C)** Representative allele plot of erythroid cells generated by ex vivo differentiation of bone marrow cells from mouse #2. The intended EPOR$^{W439*}$-edited allele was preferentially enriched during erythroid differentiation, increasing from 37.58% in the starting bone marrow cell population to 84.76% in the expanded erythroid cells. The sgRNA-binding region is indicated by the gray bar, and the red boxes highlight the intended EPOR editing site. Nucleotide substitutions relative to the reference sequence are shown in bold.

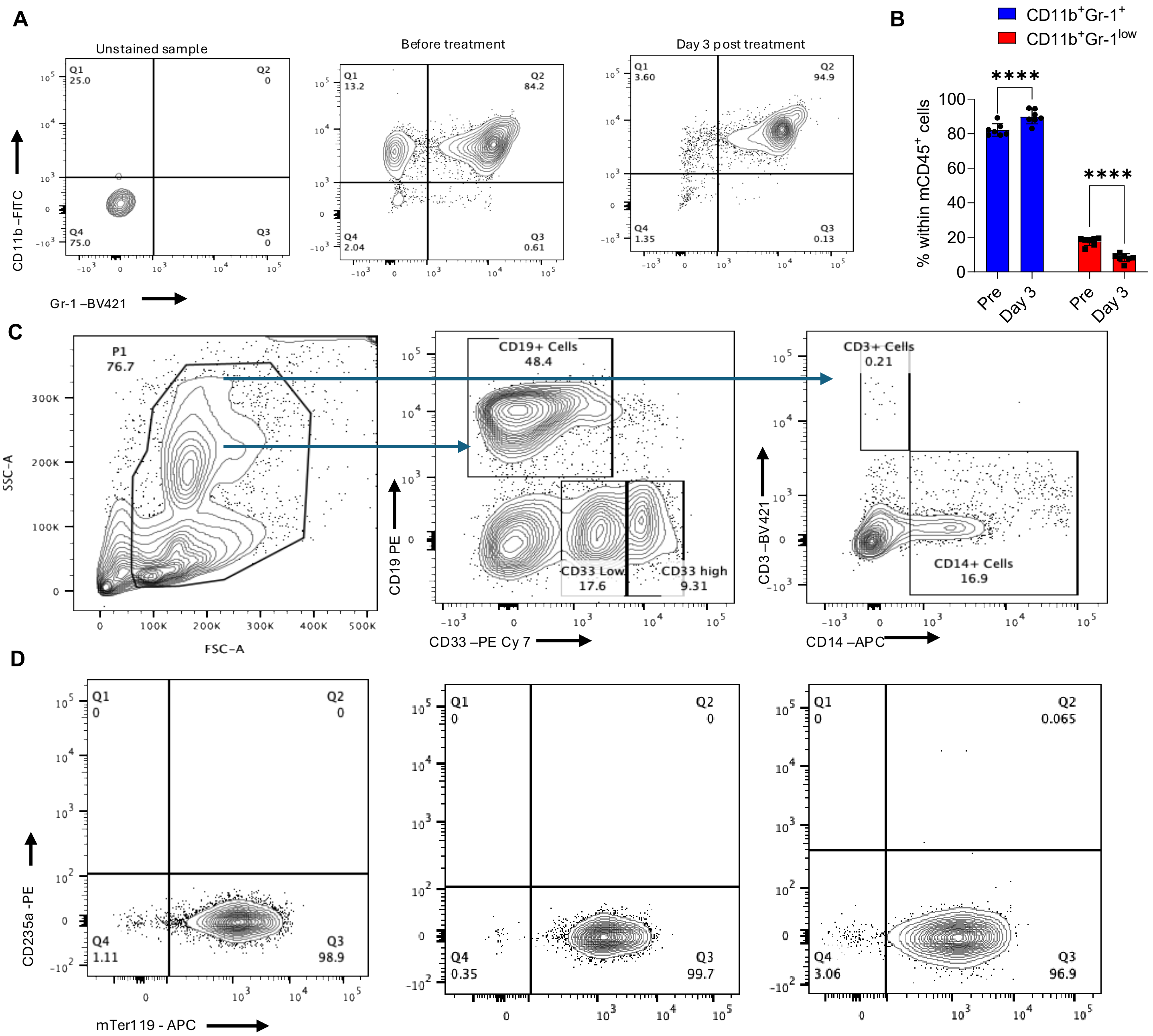


**Supplemental Figure 5. Macrophage depletion using clodronate liposomes and impact on hematopoietic and erythroid populations in transplanted NCG-X mice. (A)** Representative flow cytometry plots showing CD11b (FITC) versus Gr-1 (BV421) staining of peripheral blood cells. Unstained sample defines background. Prior to treatment, a mixed population of CD11b$^+$Gr-1$^+$ (myeloid cells) and CD11b$^+$Gr-1$^{low}$ cells, encompassing macrophages, was observed. On day 3 post-treatment with clodronate liposomes, a marked reduction in CD11b$^+$Gr-1$^{low}$ cells was evident, indicating effective depletion of macrophage/monocyte populations. **(B)** Quantification of CD11b$^+$Gr-1$^+$ (blue) and CD11b$^+$Gr-1$^{low}$ (red) populations as a percentage of CD45$^+$ cells before (Pre) and 3 days after treatment. Clodronate liposome treatment significantly reduces the CD11b$^+$Gr-1$^{low}$ population, while the CD11b$^+$Gr-1$^+$ population remains relatively unchanged. Data are shown as mean ± SEM with individual data points. ****$p < 0.0001$. **(C)** Representative flow-cytometric gating strategy used to identify major hematopoietic-lineage populations, including CD19$^+$ B cells, CD33$^{low}$ and CD33$^{high}$ myeloid cells, CD3$^+$ T cells, and CD14$^+$ monocytes. **(D)** Representative peripheral blood flow cytometry plots showing mouse Ter119 and human CD235a expression in non-humanized controls and in mice transplanted with untransduced or HDAd.EPOR$^{W439*}$-LCR-γ-GFP transduced human cells. No detectable mTer119$^-$CD235a$^+$ circulating human erythroid population was observed in the analyzed groups. Data are presented as mean ± SEM. Each dot represents an individual mouse. Statistical significance was determined using unpaired two-tailed t-test. ****$P < 0.0001$.

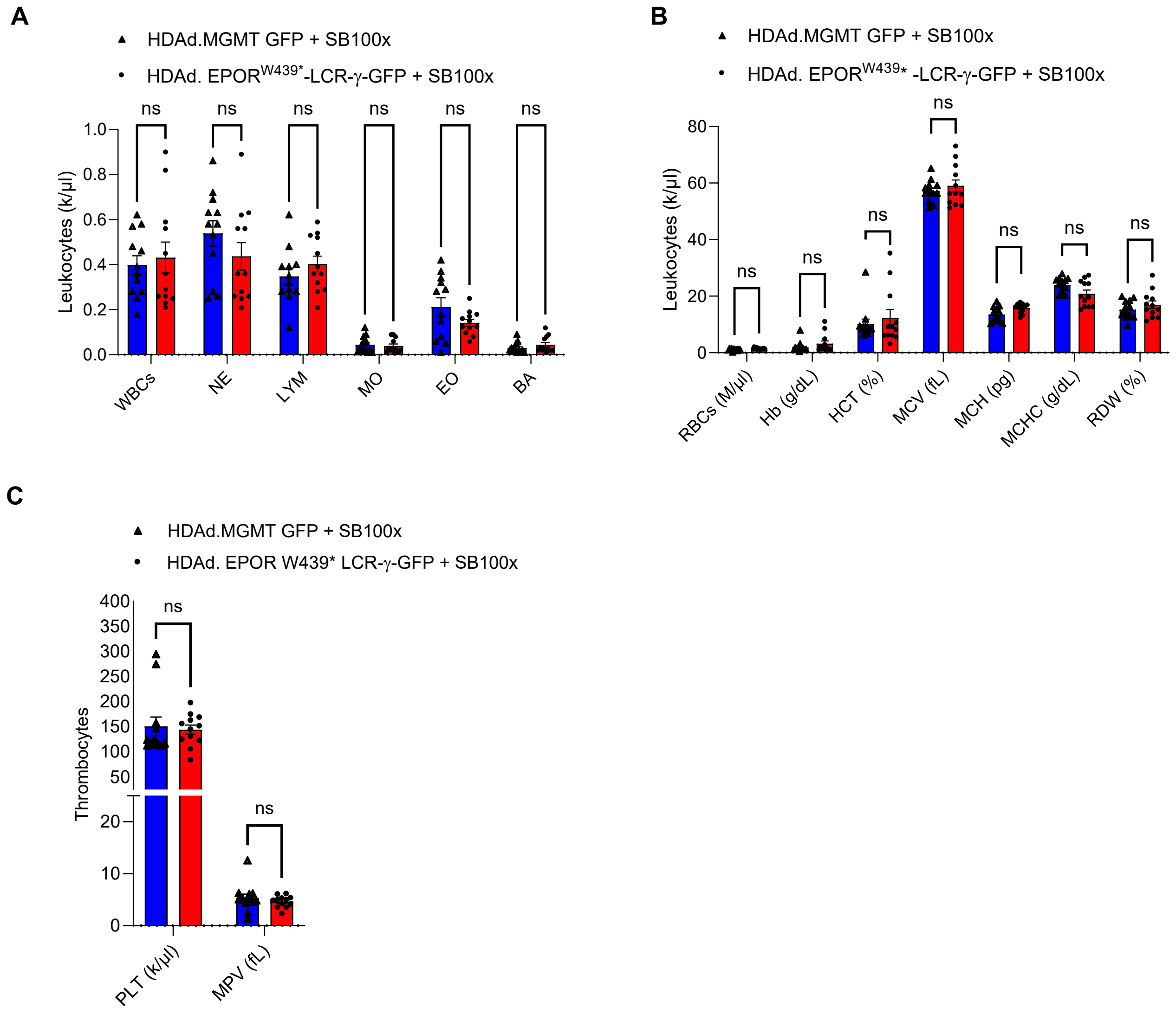


**Supplemental Figure 6. Peripheral blood hematologic parameters following HDAd transduction.** Peripheral blood was analyzed in mice treated with the control vector HDAd.MGMT/GFP plus HDAd.SB100x or with HDAd.EPOR$^{W439*}$-LCR-γ-GFP plus HDAd.SB100x. **(A)** Absolute leukocyte counts, including total white blood cells (WBCs), neutrophils (NE), lymphocytes (LYM), monocytes (MO), eosinophils (EO), and basophils (BA). **(B)** Erythrocyte parameters, including red blood cell count (RBC), hemoglobin concentration (Hb), hematocrit (HCT), mean corpuscular volume (MCV), mean corpuscular hemoglobin (MCH), mean corpuscular hemoglobin concentration (MCHC), and red cell distribution width (RDW). **(C)** Platelet count (PLT) and mean platelet volume (MPV). Bars represent the mean ± SEM, and individual points represent individual mice.

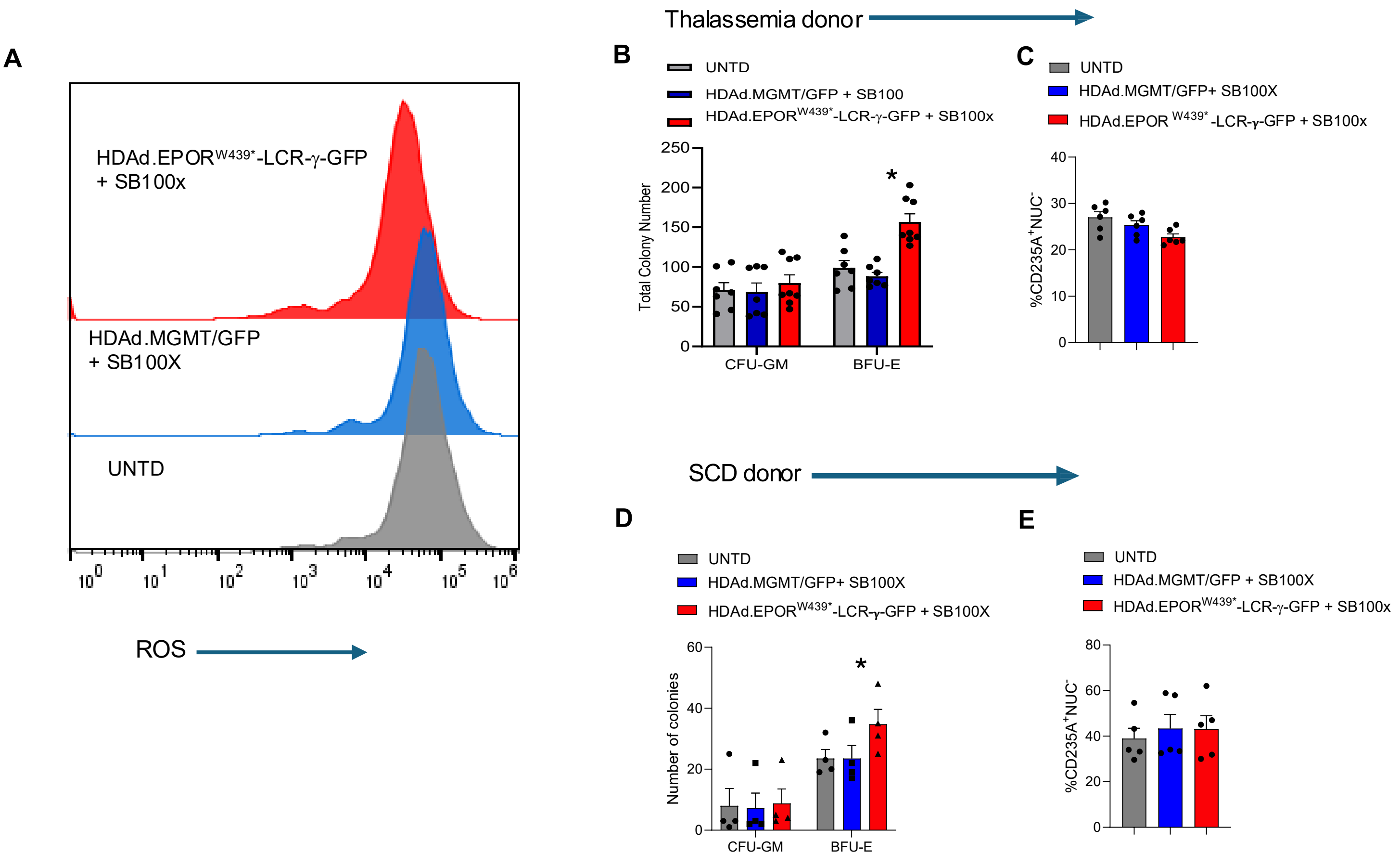


**Supplemental Figure 7. EPOR$^{W433*}$ editing combined with γ-globin gene addition supports erythroid differentiation and maturation in β-thalassemia- and SCD–derived CD34$^{+}$ cells. (A to C)** Studies with β-thalassemia patient-derived CD34$^{+}$ cells. **(A)** Representative flow-cytometry plots showing intracellular reactive oxygen species (ROS) levels in erythroid cells. **(B)** Distribution of progenitor colonies. **(C)** Frequency of enucleated CD235a$^{+}$ erythroid cells (CD235a$^{+}$NUC$^{-}$). **(D to E)** Studies with SCD patient-derived CD34$^{+}$ cells. **(D)** Numbers of granulocyte–macrophage colony-forming units (CFU-GM) and erythroid burst-forming units (BFU-E). **(E)** Frequency of enucleated CD235a$^{+}$ erythroid. Gray represents untransduced cells, blue represents HDAd.MGMT/GFP + HDAd.SB100x, and red represents HDAd.EPOR$^{W439*}$-LCR-γ-GFP + HDAd.SB100x. Bars show mean ± SEM with individual mice indicated.

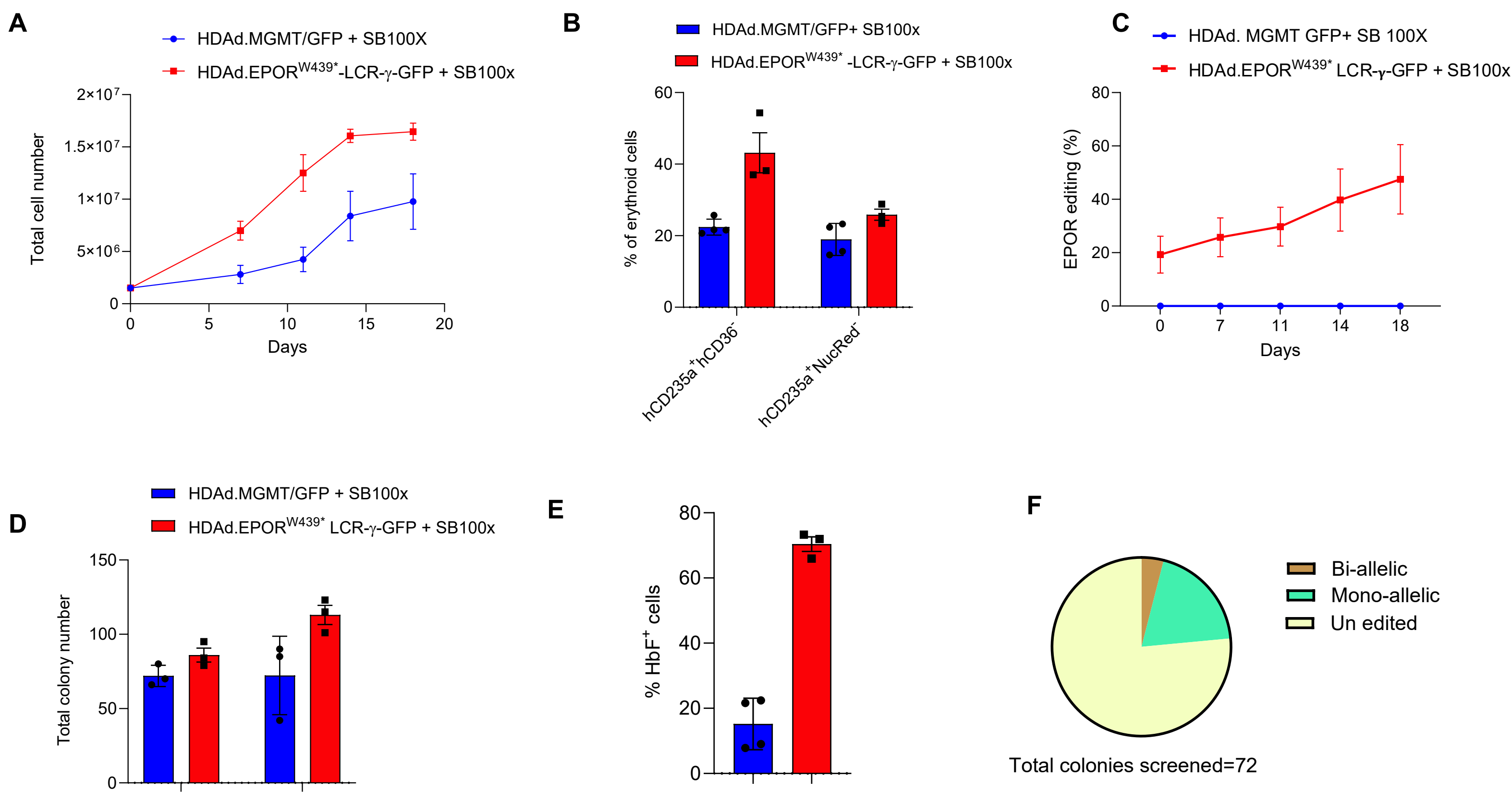


**Supplemental Figure 8. In vivo EPOR-W439* editing combined with γ-globin gene addition produces higher numbers of HbF⁺ erythroid progeny from bone marrow cells isolated from humanized mice. (A)** Total cell numbers during ex vivo erythroid differentiation of cells isolated from bone marrow of mice treated with HDAd.MGMT/GFP + HDAd.SB100x or HDAd.EPOR$^{W439*}$-LCR-γ-GFP + HDAd.SB100x. **(B)** Frequencies of erythroid-cell populations defined by human CD235a, CD36, and nuclear-dye staining following ex vivo differentiation. **(C)** Longitudinal analysis of EPOR$^{W439*}$ editing during erythroid differentiation, demonstrating progressive enrichment of edited cells. **(D)** Total numbers of hematopoietic colonies generated from cells isolated from the indicated treatment groups. **(E)** Frequencies of HbF-positive cells following erythroid differentiation. **(F)** Clonal analysis of EPOR editing in individual hematopoietic colonies derived from EPOR$^{W439*}$-edited CD34$^{+}$ cells. Colonies were classified as bi-allelic, mono-allelic, or unedited based on EPOR genotype; a total of 72 colonies were analyzed. Individual points represent independent animals or cultures, and data are presented as mean ± SEM.

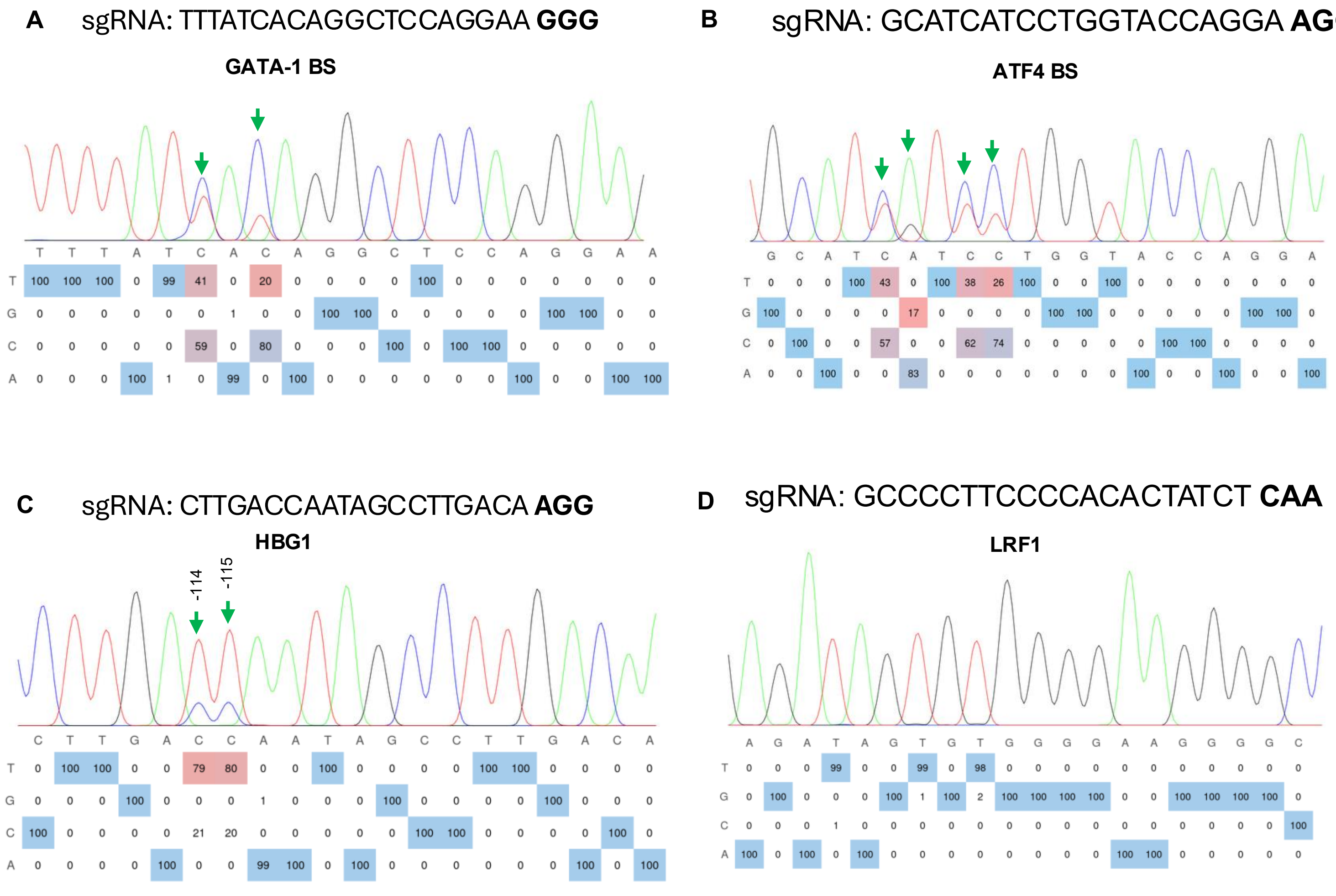


**Supplemental Figure 9. Sequence validation of base editing at HbF-regulatory target sites.** Representative Sanger-sequencing chromatograms and nucleotide-frequency analyses of cells edited at the GATA1-binding site **(A)**, ATF4-binding site **(B)**, HBG promoter sgRNA-2 target site **(C)**, and LRF1 target site **(D)**. Green arrows indicate the intended nucleotides detected within the base-editor activity window. In panel C, the targeted HBG promoter nucleotides at positions −114 and −115 relative to the transcription start site are indicated. The tables below each chromatogram show the percentage of A, C, G, and T detected at each nucleotide position; mixed nucleotide peaks reflect edited and unedited alleles within the analyzed cell population.

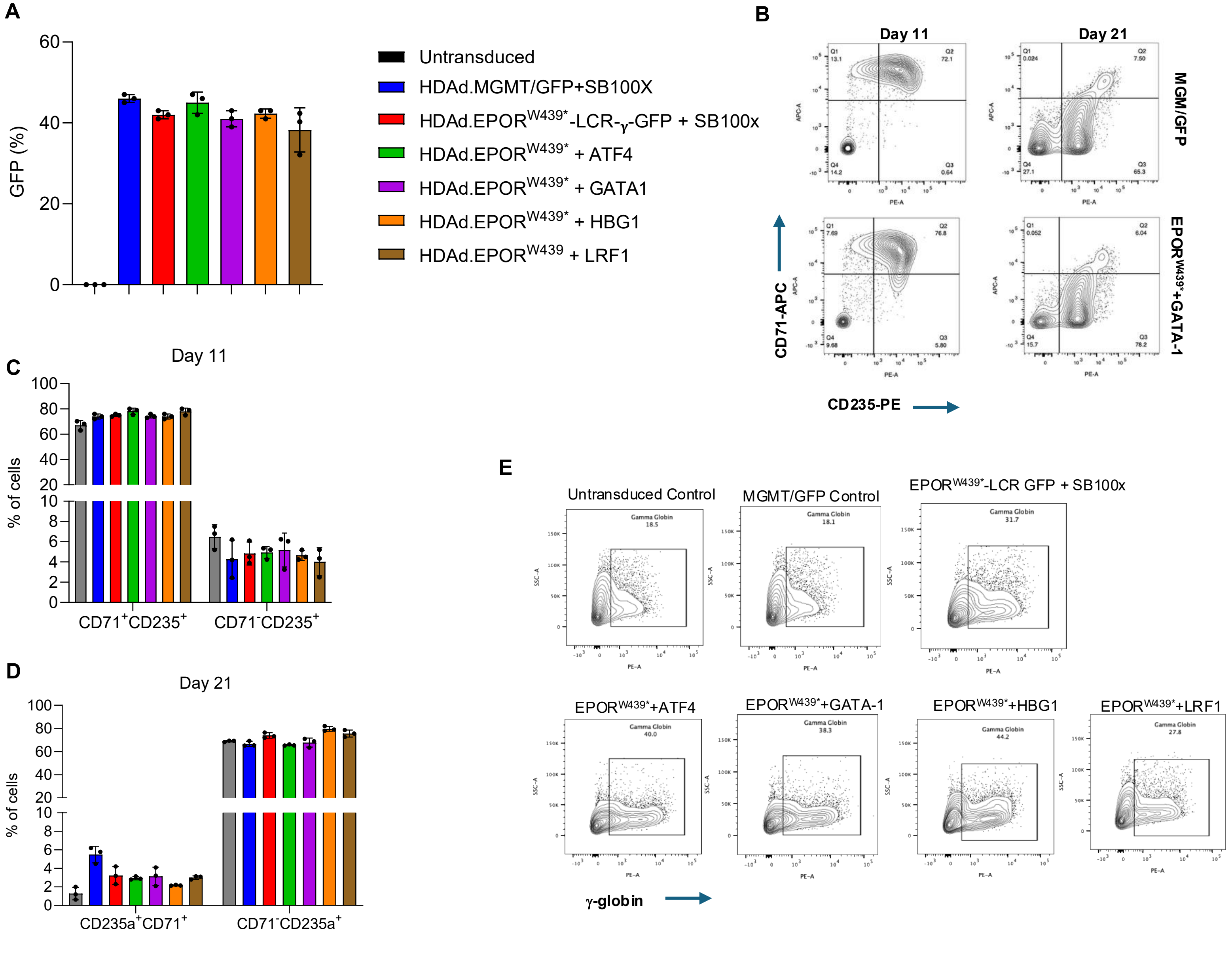


**Supplemental Figure 10. HDAd-mediated dual base editing preserves erythroid differentiation and reactivates HbF expression in human HSPCs. (A)** Transduction efficiency of human CD34$^+$ cells from healthy donors following treatment with the indicated HDAd vector combinations, assessed by the percentage of GFP-positive cells. Untransduced cells served as a negative control. **(B)** Representative flow-cytometry plots showing erythroid differentiation on days 11 and 21 in cells. CD34$^+$ cells were transduced (4000 vp/cell) with the indicated HDAd vectors and subjected to three-phase erythroid differentiation. Cells were collected on Day 11 and Day 21 to assess differentiation potential by CD71 and CD235a expression. **(C)** Representative intracellular flow-cytometry plots showing γ-globin reactivation in untransduced cells and cells treated with HDAd.MGMT/GFP control, HDAd.EPOR$^{W439*}$-LCR-γ-GFP plus HDAd.SB100x, or dual-editing HDAd vectors targeting EPOR together with ATF4, GATA1, HBG1 promoter, or LRF1. The gated regions indicate HbF-positive cells, with the corresponding percentages shown in each plot. Data in panel A are presented as mean ± SEM from three independent donor, with individual measurements shown.

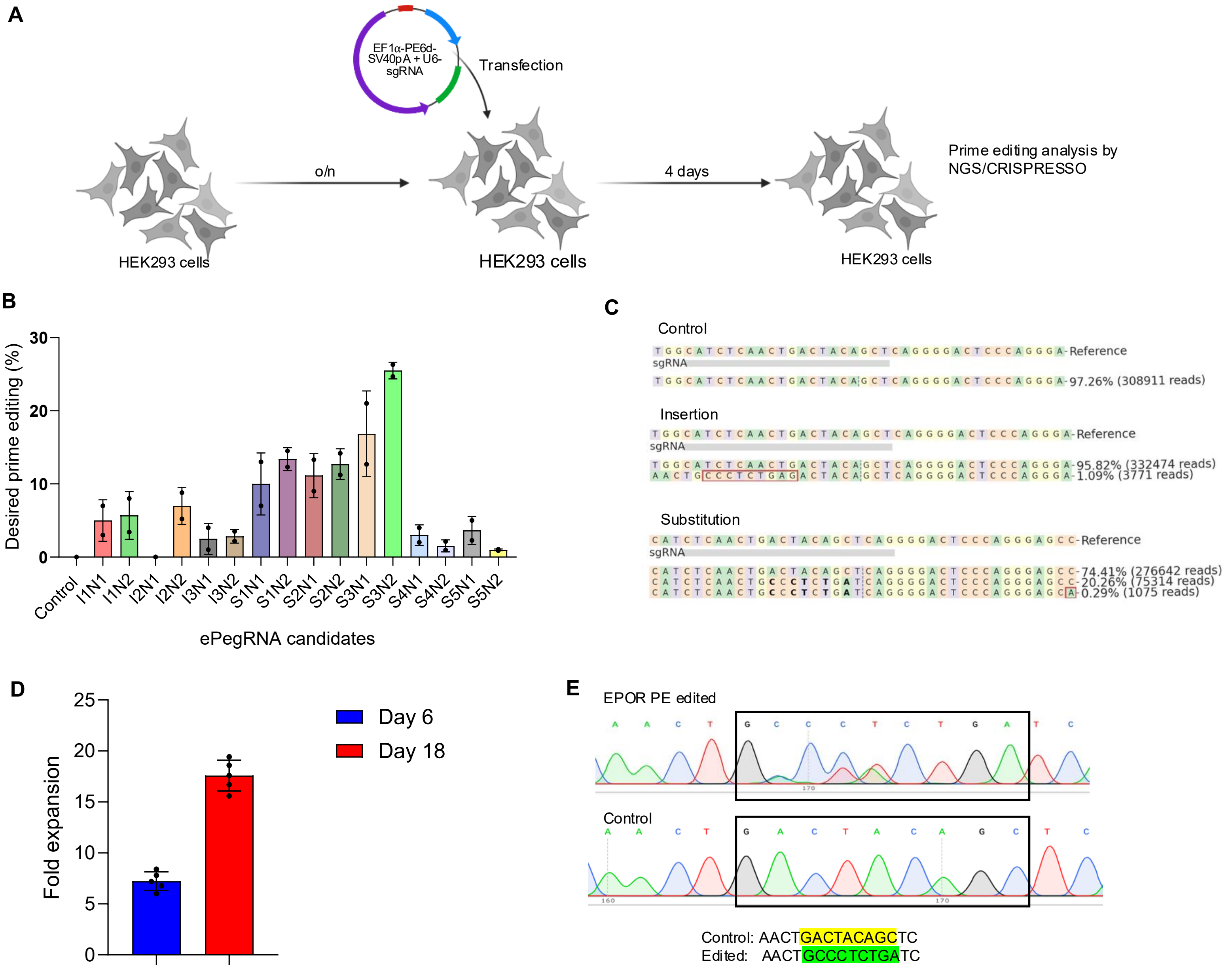


**Supplemental Figure 11. Optimization and validation of prime editing at the EPOR target site. (A)** Experimental workflow for screening prime-editing guide RNA combinations. HEK293 cells were transfected with plasmids encoding the PE6d prime editor and the U6-driven pegRNA/nickRNA combinations. Four days after transfection, genomic DNA was collected and editing outcomes were quantified by targeted next-generation sequencing and CRISPResso analysis. **(B)** Frequencies of the desired EPOR prime-editing outcome obtained with the indicated guide RNA candidates. Individual points represent replicate measurements, and bars show mean ± SEM. **(C)** Representative allele plots showing the unedited control sequence and the predominant insertion- and substitution-containing alleles detected after prime editing. The frequencies and read numbers of the major alleles are indicated. **(D)** Fold expansion of HUDEP-2 cells harboring the selected EPOR edit on days 6 and 18 of culture, demonstrating progressive enrichment of the edited population. HUDEP-2 cells were transduced with HDAd.Dual PE vector at MOI 1000 vp/cell. After 4 days, cells were counted, and an equal number of control and EPOR-prime-edited cells were expanded for 18 days. Fold expansion of EPOR-prime edited cells calculated relative to unedited control. **(E)** Representative Sanger-sequencing chromatograms confirming the intended EPOR sequence modification in prime-edited cells compared with unedited control cells. The corresponding control and edited sequences are shown below the chromatograms, with the modified nucleotides highlighted. HUDEP-2 cells at Day 18 were collected from control and EPOR-prime edited conditions and editing was analyzed using Sanger sequencing. Data are presented as mean ± SEM, with individual measurements shown.

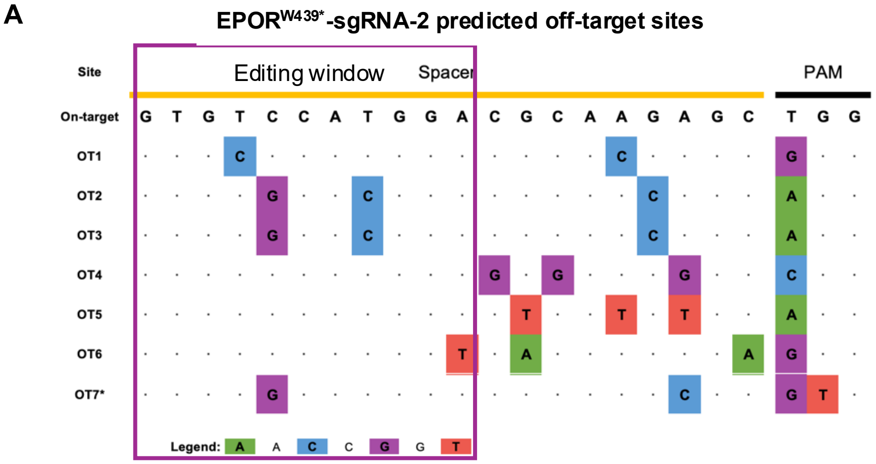


| | | | |
|---|---|---|---|
| OT1 | Chr1:37484603-37484625 | OT5 | Chr19:46344555-46344577 |
| OT2 | Chr2:110079662-110079684 | OT6 | Chr14:98949905-98949927 |
| OT3 | Chr2:110291692-110291714 | OT7 | Chr12:132558472-132558494 |
| OT4 | Chr22:16387656-16387678 | | |

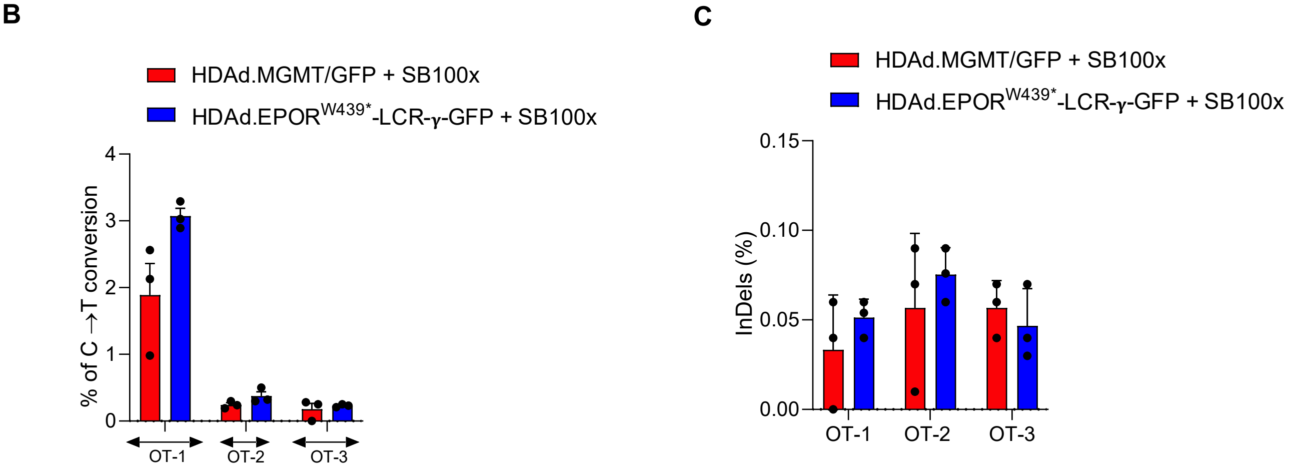


**Supplemental Figure 12. Targeted evaluation of predicted off-target editing by the EPOR$^{W439*}$ sgRNA-2. (A)** Sequence alignment of the EPOR$^{W439*}$ sgRNA-2 on-target site and seven candidate off-target loci identified by COSMID. The 20-nt spacer and PAM are indicated, and the analyzed base-editing window is highlighted. Dots denote nucleotides identical to the on-target sequence, whereas colored letters indicate mismatched nucleotides. OT2 and OT3 contain the same spacer/PAM sequence but map to distinct genomic loci. OT7, marked with an asterisk, contains a noncanonical GTG PAM. The hg38 genomic coordinates of each candidate site are shown below the alignment. **(B)** Frequencies of C-to-T conversion at cytosines located within the analyzed editing window at OT1, OT2, and OT3. The evaluated cytosines were located at spacer position 4 for OT1 and position 8 for OT2 and OT3. **(C)** Indel frequencies detected at OT1–OT3. Human CD34$^{+}$ cells treated with the control HDAd.MGMT/GFP plus HDAd.SB100x vector combination were compared with cells treated with HDAd.EPOR$^{W439*}$-LCR-γ-GFP plus HDAd.SB100x. Bars represent mean ± SEM, and individual points represent independent donor samples.

A

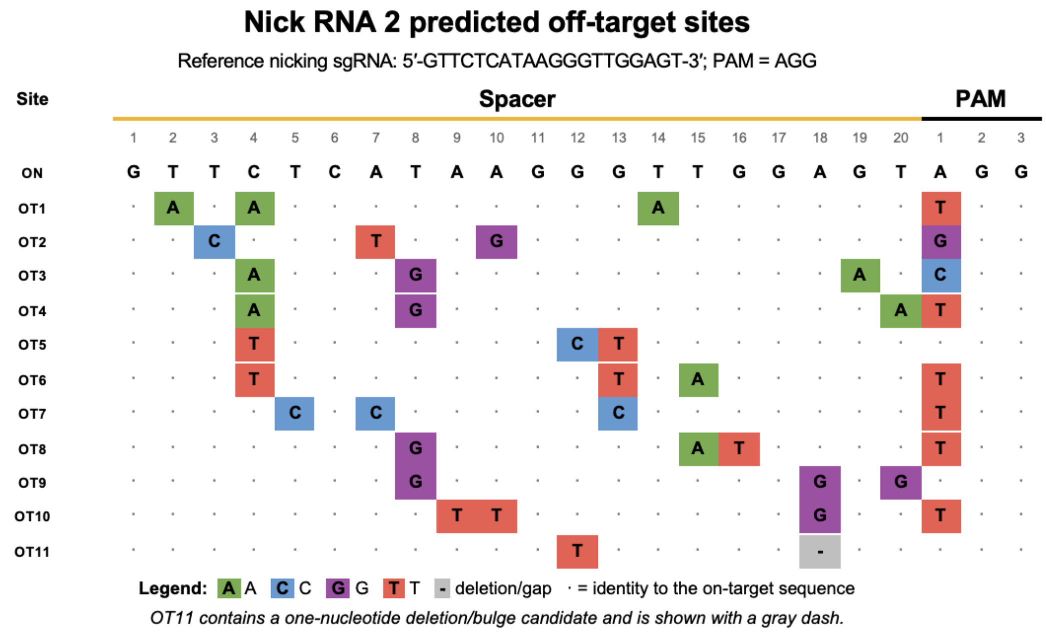


| Site | Genomic coordinate (hg38) | Site | Genomic coordinate (hg38) |
|---|---|---|---|
| OT1 | Chr13:97781758-97781780 | OT7 | Chr5:137459352-137459374 |
| OT2 | Chr11:35567158-35567180 | OT8 | Chr3:6100389-6100411 |
| OT3 | Chr3:127257137-127257159 | OT9 | Chr15:52851681-52851703 |
| OT4 | Chr2:85326916-85326938 | OT10 | Chr6:68810322-68810344 |
| OT5 | ChrX:47983887-47983909 | OT11 | Chr2:16716743-16716764 |
| OT6 | Chr16:68503262-68503284 | | |

Dots indicate identity to the on-target sequence. Colored boxes indicate nucleotide mismatches.

B

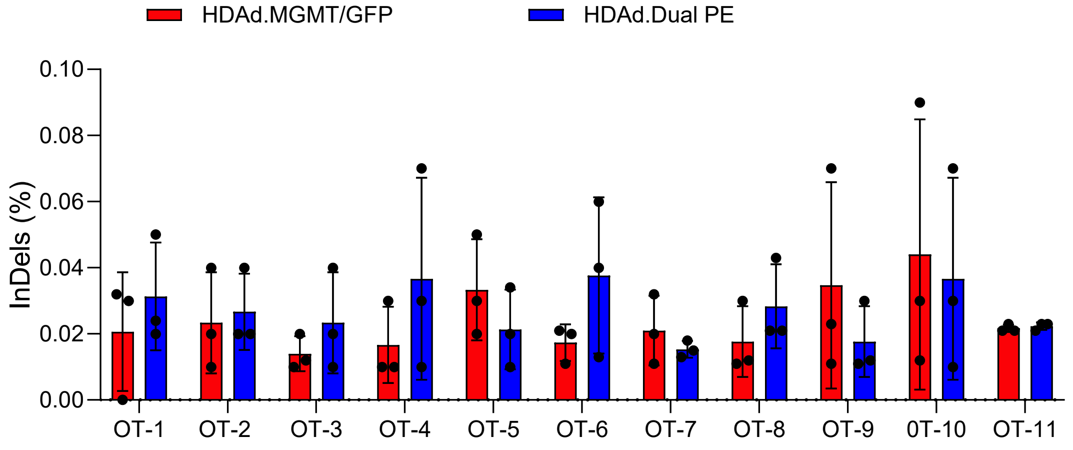


**Supplemental Figure 13. Targeted analysis of predicted off-target sites for the prime-editing nicking sgRNA. (A)** Alignment of the EPOR nick RNA 2 spacer sequence (5′-GTTCTCATAAGGGTTGGAGT-3′; PAM, AGG) with 11 predicted genomic off-target sites identified using COSMID. Dots indicate nucleotide identity with the on-target spacer, colored boxes indicate mismatched nucleotides, and the gray dash denotes the one-nucleotide deletion/bulge predicted at OT11. The corresponding genomic coordinates are shown according to the GRCh38/hg38 assembly. **(B)** Indel frequencies at the 11 predicted off-target sites were quantified by targeted amplicon sequencing in cells transduced with the control vector HDAd.MGMT or the dual prime-editing vector HDAd.Dual-PE. Bars represent the mean ± SEM, and individual points represent independent donors (n = 3). Indel frequencies remained low across all analyzed sites.

**A**

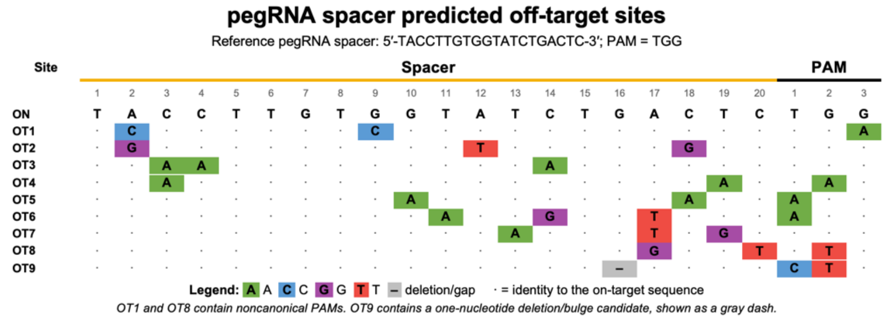


| Site | Genomic coordinate (hg38) | Site | Genomic coordinate (hg38) |
|---|---|---|---|
| OT1 | ChrX:118726580-118726602 | OT6 | Chr2:178771052-178771074 |
| OT2 | Chr1:44316376-44316398 | OT7 | Chr2:135817740-135817762 |
| OT3 | Chr20:17354641-17354663 | OT8 | ChrX:82420478-82420500 |
| OT4 | Chr1:71723858-71723880 | OT9 | Chr5:161934844-161934865 |
| OT5 | Chr3:65128405-65128427 | | |

Dots indicate identity to the on-target sequence. Colored boxes indicate nucleotide mismatches.

**B**

**C**

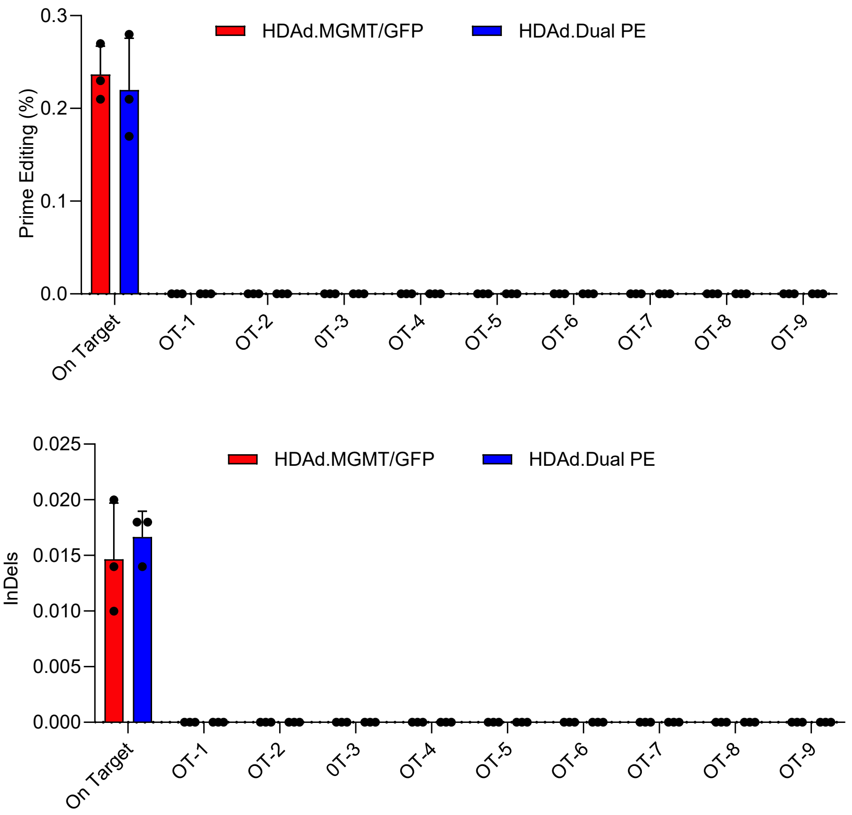


**Supplemental Figure 14. Targeted analysis of predicted off-target sites for the EPOR pegRNA spacer. (A)** Alignment of the EPOR pegRNA spacer sequence (5′-TACCTTGTGGTATCTGACTC-3′; PAM, TGG) with nine predicted genomic off-target sites identified using COSMID. Dots indicate nucleotide identity with the on-target sequence, colored boxes indicate nucleotide mismatches, and the gray dash denotes the predicted one-nucleotide deletion/bulge at OT9. OT1 and OT8 contain noncanonical PAM sequences. Genomic coordinates are shown according to the GRCh38/hg38 human genome assembly. **(B)** Frequencies of the exact programmed prime-editing outcome at the on-target locus and nine predicted pegRNA off-target sites were quantified by targeted amplicon sequencing in cells transduced with the control vector HDAd.MGMT/GFP or the dual prime-editing vector HDAd.Dual-PE. **(C)** Indel frequencies at the same loci. Bars represent the mean ± SEM, and individual points represent independent donors (n = 3). No detectable prime-editing products or indels above background were observed at the predicted off-target sites.
